# NSD2 deficiency disrupts the epigenetic landscape of cochlear hair cells to enhance ECM expression and to perturb cochlear hair cell function

**DOI:** 10.64898/2026.09.21.753379

**Authors:** Ziyi Wang, Yue Xu, Xiaojie Ma, Dehuan Wang, Wenxin Feng, Ningyuan Liu, Hanyu Rao, Wei Zhang, Rebiguli Aji, Ziwen Yu, Jiahe Li, Guoning Yu, Yazhi Xing, Wei-Qiang Gao, Li Li

## Abstract

Mammalian cochlear hair cells (HC) are crucial for hearing, and histone modifications may play an important role in HC maturation and functions. However, whether and how NSD2, a histone H3 lysine 36 (H3K36) dimethyltransferase, affects cochlear HC maturation remains unclear. Here, we established an HC-specific Nsd2 knockout mouse model and discovered that the loss of NSD2 results in severe structural defects in stereociliary bundles and profound hearing impairment. Integrated multi-omics profiling revealed that NSD2 deficiency leads to a decreased expression of H3K36me2, which in turn alters genome-wide chromatin accessibility and gene expression. Mechanistically, NSD2 deficiency triggers the aberrant upregulation of extracellular matrix (ECM) genes at the transcriptional level, causing excessive collagen accumulation and compromising tissue structure. Crucially, ECM intervention can mitigate NSD2-deficiency mediated hearing damage. Our study reveals an epigenetic mechanism by which NSD2 regulates cochlear HC maturation through inhibition of chromatin remodeling-induced ECM accumulation, and provides a potential new therapeutic direction for hearing impairment.

## Introduction

Hearing impairment represents one of the most prevalent sensory dysfunctions in humans, severely compromising inter-individual communication and cognitive capacities. Clinically, hearing loss is classified into conductive, sensorineural, and mixed types, driven by an array of etiological factors including genetic mutations, noise exposure, ototoxic drugs, infectious diseases, and aging^1^. Although hearing aids and cochlear implants remain the primary therapeutic interventions for sensorineural hearing loss^2^, the surgical trauma of implantation can occasionally precipitate adverse cochlear fibrosis and ossification. Therefore, unraveling the molecular mechanisms underlying hearing impairment is a pressing issue.

In the mammalian auditory system, hair cells (HCs) are capable of converting the mechanical vibrations of external sounds into intracellular electrochemical signals^3–5^. They are classified as inner and outer hair cells, and synapse with the peripheral processes of spiral ganglion neurons (SGNs) to transmit auditory signals to the central nervous system. HC development is primarily linked to genes including *Math1*^6^ (*Atoh1*), *Hes1*^7^, and multiple signaling pathways^8–11^, which are important for the proliferation and differentiation of HC progenitors.

Auditory function relies heavily on the intricate interplay between cochlear HCs and the extracellular matrix (ECM) within the organ of Corti^12^. Composed primarily of collagens and specialized cochlear proteins, the inner ear ECM forms the structural scaffolds of the basilar membrane (BM) and tectorial membrane (TM) ^13–15^. Recent evidence highlights that the ECM structurally directs cytoskeleton remodeling and guides cell differentiation trajectories during development^16^. For instance, mice deficient in the collagenous fiber Emilin-2 exhibit impaired auditory thresholds, demonstrating that ECM defects disrupt BM biomechanical properties and compromise hearing^17^. Furthermore, mutations in critical collagen genes (e.g., *Col2a1, Col4a3, Col4a4, Col4a5, Col9a1, Col11a1*, and *Col11a2*) directly cause syndromic hearing impairments^18^. Collectively, precise fine-tuning of cochlear ECM remodeling is indispensable for maintaining proper biomechanical and functional properties in the cochlea.

On the other hand, epigenetic modifications play an equally essential role in hearing maturation ^19^. Accumulating evidence indicates that histone modifications may also play a significant role in the proliferation, differentiation, and fate determination of inner ear stem cells^20^. For instance, the differentiation of inner ear neurons relies on the transcriptional enhancement of Neurog1 facilitated by histone acetylation^21^. Meanwhile, expression of *Atoh1*, a crucial gene for the development of inner ear HC, is regulated by histone acetylation. Inhibition of histone acetyltransferase activity downregulates *Atoh1* and induces defects in HC development^22^. In terms of methylation, the histone demethylase LSD1 restricts the accumulation of H3K4me and H3K4me2 to maintain the pluripotency of ear progenitors^23,24^. The regulatory changes in histone acetylation and methylation suggest that histone modifications control gene expression in a developmentally stage-specific manner, playing a pivotal role in cochlear development and the differentiation of inner ear HCs.

NSD2, alternatively known as WHSC1 or MMSET, belongs to the nuclear receptor-binding SET domain (NSD) protein family, which exhibits high evolutionary conservation between humans and mice^25–27^. It is mainly responsible for catalyzing the monomethylation and dimethylation of histone H3K36^28^. The NSD histone methyltransferase family also includes NSD1 (KMT3B) and NSD3 (WHSC1L1). The catalytic activity of NSD proteins depends on their binding to nucleosomes, accompanied by a dynamic transition between active and inactive conformations. Loss-of-function and mis-sense mutations of NSD2 lead to reduced methylation activity, which is associated with distinct developmental phenotypes such as sensorineural deafness in Wolf-Hirschhorn syndrome^29–32^. Notably, an NSD2-deficient model exhibited malformed apical stereociliary bundles of sensory HCs and defective neural innervation.^32^ However, the role and molecular mechanisms of NSD2 during cochlear HC maturation have not been explored yet.

Here, we report a link between NSD2-deficiency and aberrant ECM accumulation in cochlear HC maturation. To elucidate the mechanism of hearing impairment caused by NSD2 deficiency, we generated HC-specific *Nsd2* knockout (*Nsd2^Atoh^*^1^*^-KO^*) mice. Our results demonstrate that the loss of NSD2 disrupts the H3K36me2-associated epigenetic landscape in cochlear HCs, which subsequently triggers the aberrant upregulation of ECM genes. Collectively, these findings fill the gap between H3K36me2 epigenetic regulation and the hearing impairment caused by NSD2 deficiency.

## Results

### NSD2 expression is associated with early HC maturation

To explore the relationship between NSD2 and cochlear maturation, we analyzed the existing public sequencing datasets of mouse (NCBI Gene Expression Omnibus: GSE53863 and GSE60019). Transcriptional analysis of the GSE53863 dataset revealed that Nsd2 expression was highest in the apical turn of the cochlea compared to the middle and basal turns in 6-week-old mice (Fig. 1A), reflecting a molecular trace of the developmental gradient across the cochlear apex-base axis. To correlate this spatial pattern with early maturation, we next analyzed Nsd2 expression across critical pre- and postnatal stages. Analysis of the GSE60019 dataset revealed stage-dependent variations in Nsd2 expression, suggesting its dynamic modulation during cochlear hair cell maturation (Fig. S1A). We performed RT-qPCR using murine organs of Corti across several developmental time points (Fig. 1B). The data showed that *Nsd2* expression exhibited temporal fluctuations across different stages, suggesting its dynamic involvement in the development and maturation of the Organ of Corti (Fig. 1B).

**Fig. 1.**
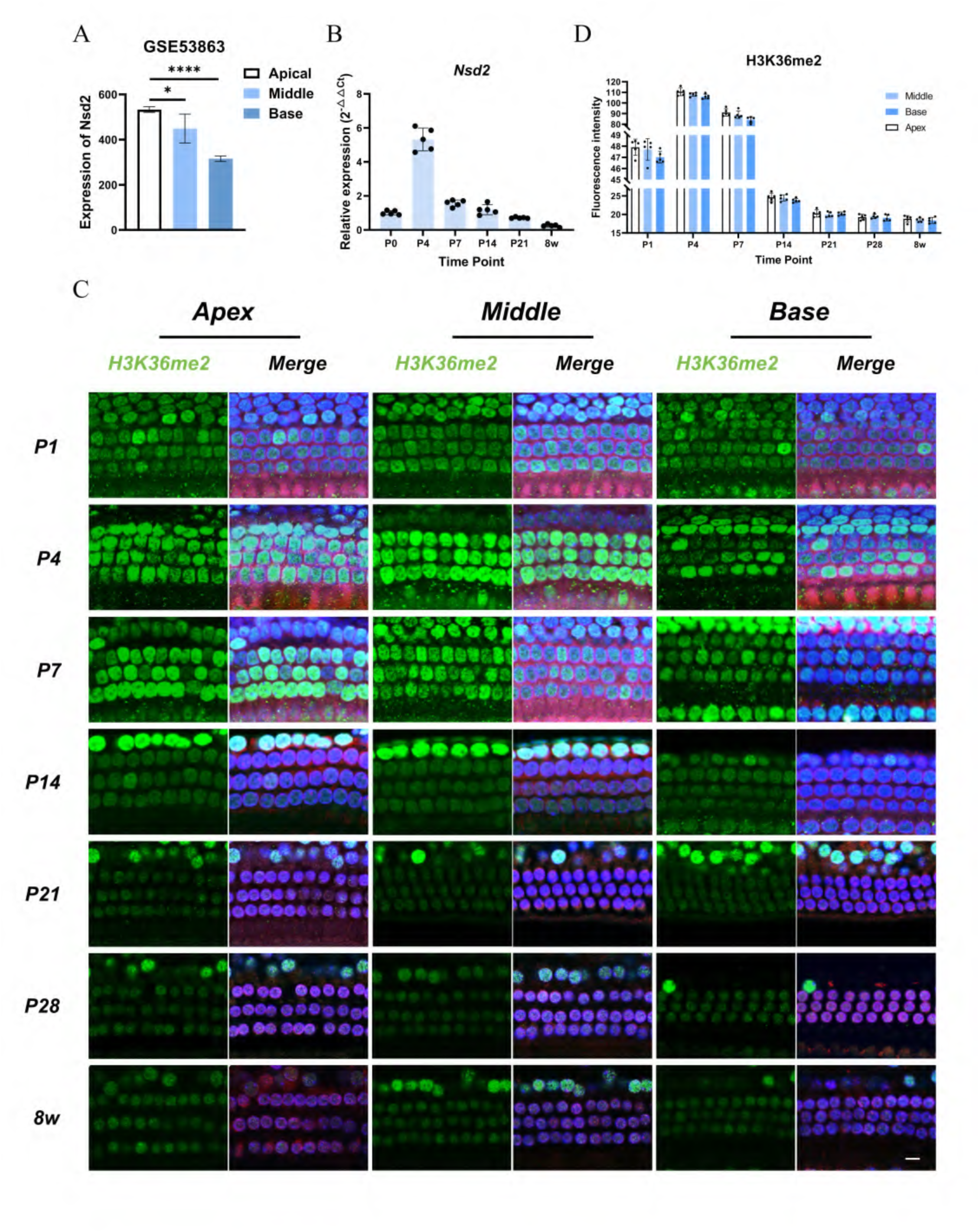
NSD2 expression is associated with early HC maturation. **(A)** Normalized mRNA expression levels of Nsd2 across the apical, middle, and basal turns of normal cochlear tissues, retrieved from the public transcriptomic dataset GSE53863. The data demonstrate a distinct base to apex spatial gradient, with Nsd2 transcripts being significantly enriched within the less-differentiated apical region. Data are presented as mean ±SEM. *P < 0.05, and ****P < 0.0001 **(B)** Temporal mRNA expression profile of Nsd2 in the mouse auditory epithelium during postnatal maturation. Quantitative RT-qPCR analysis was performed using freshly dissected whole organ of Corti bulk tissues from wild-type C57BL/6J mice across sequential developmental stages, ranging from postnatal day 0 (P0) to adulthood (8w). Relative expression levels were calculated via the 2^-ΔΔCT^ method and normalized to Gapdh enrichment. Individual data points represent independent biological replicates (n = 6 per group), and data are plotted as mean ± S.D. One-way ANOVA with Tukey’s post hoc test. **(C)** Spatiotemporal distribution of H3K36me2 enrichment along the cochlear tonotopic axis during postnatal maturation. Representative whole-mount immunofluorescence surface images display the sensory epithelium across the apical (Apex), middle (Middle), and basal (Base) turns of wild-type C57BL/6J mice from postnatal day 1 (P1) to adulthood (8w). Left panels illustrate independent H3K36me2 immunoreactivity (green), and right panels illustrate merged channels (Merge) consisting of the cell-type-specific HC marker Myo7a (red), H3K36me2 (green), and DAPI (blue). To guarantee rigorous quantitative comparison and eliminate depth artifacts, all laser scanning confocal micrographs were acquired under identical laser and gain parameters at a single horizontal optical plane stabilized at the cell body level. The distinct cell-specific intensity shifts validate a developmentally regulated epigenetic reprogramming wave within the HC population. Images are representative of n=4 independent cochlear preparations per developmental stage. Scale bar = 15 μm. **(D)** Quantitative analysis of H3K36me2 fluorescence intensity shifts within the maturation timeline. Semi-quantitative fluorescence intensity was measured from laser scanning confocal images within the defined apical (Apex), middle (Middle), and basal (Base) turns across sequential postnatal stages through adulthood (8w). For cell-specific measurement, Regions of Interest (ROIs) were circumscribed around individual somatic boundaries defined by the canonical HC marker Myo7a using ImageJ software. To guarantee technical accuracy and eliminate spatial bias, the baseline value for each individual cell was calculated as the grand mean of ROIs measured across 3 to 4 continuous single horizontal optical planes captured at the exact same spatial location. Each plotted data point represents an independent biological replicate (n = 5 per group). Data are plotted as mean ± S.D. along a split Y-axis configuration to capture high-resolution signal amplitude variances. Two-way ANOVA with Tukey’s post hoc test.

Given that NSD2 acts as a primary histone methyltransferase targeting H3K36me2, we next performed immunofluorescence staining to quantify H3K36me2 intensity from HC across the apical, middle, and basal turns of the cochlea (Fig. 1C and statistical analysis in Fig. 1D). Quantitative analysis revealed that H3K36me2 levels synchronously surged and peaked at P4 in all three turns, followed by a progressive decline from P7 onwards, stabilizing at a minimum by P21 (Fig. 1D). Intriguingly, across almost all examined stages, H3K36me2 fluorescence intensity was consistently higher in the apical turn compared to the middle and basal turns (Fig. 1D). These temporal kinetics demonstrate a tight coupling between *Nsd2* transcription and its mediated epigenetic modifications, while the persistent apex-base spatial gradient further reinforces the spatial asymmetry observed during HC maturation. These data collectively demonstrate that NSD2 expression correlates with early HC maturation, suggesting that its mediated H3K36me2 modification is involved in this developmental process.

### NSD2 deletion impairs mouse hearing

To determine whether NSD2 influences HC maturation and auditory function, we established a mouse model with HC-specific knockout of NSD2 (*Nsd2^Atoh^*^1^*^-KO^* mice) based on genotyping (Fig.S1 B and C) using the *Atoh1^Cre^* mouse ^33^ and *Nsd2^f/f^* mouse ^34^. The *Nsd2^f/f^* mouse was gifted by Prof. Jun Qin from Chinese Academy of Sciences. And the *Atoh1^Cre^*mouse used in this study was purchased from the Jackson Laboratory (Strain #: 011104) ^35^. Immunofluorescence staining demonstrated a profound decrease in both NSD2 and H3K36me2 expression specifically within the *Nsd2^Atoh1-KO^* cochlear HCs (Fig. 2 A and B). These data confirmed the successful knockout of NSD2 in C57BL/6 mice, providing a genetic model to evaluate its role in hearing impairment.

**Fig. 2.**
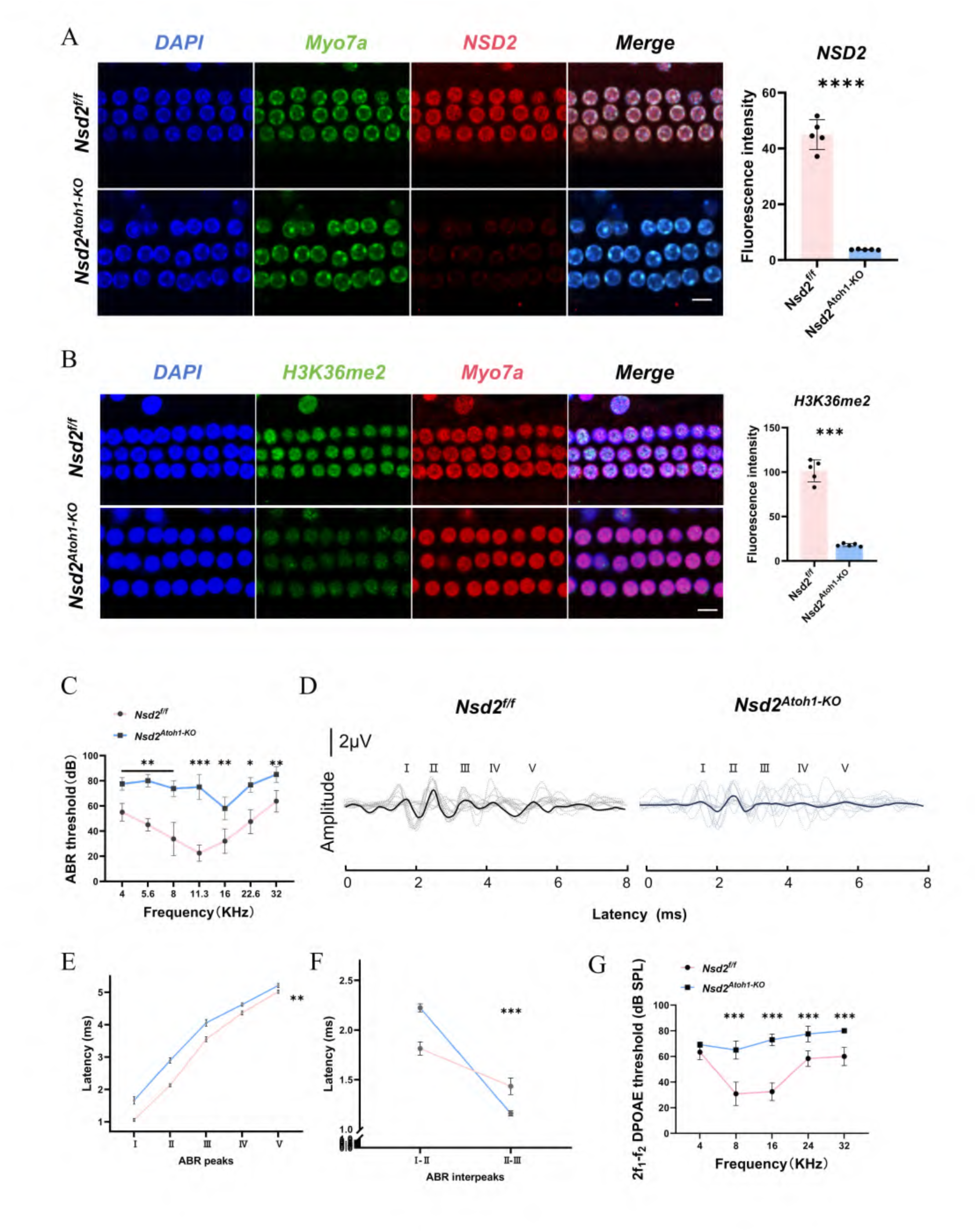
NSD2 deletion impairs mouse hearing. **(A)** Representative whole-mount immunofluorescence surface images and corresponding semi-quantitative analysis of NSD2 expression in the sensory epithelium of adult *Nsd2^f/f^* and *Nsd2^Atoh1-KO^* mice. Confocal panels illustrate nuclear counterstaining via DAPI (blue), the canonical HC lineage marker Myo7a (green), NSD2 protein labeling (red), and the merged channels (Merge). Knockout mice exhibit a profound and selective loss of NSD2 signal within the HC rows. To ensure technical reproducibility, Regions of Interest (ROIs) were circumscribed around individual cell somas delineated by Myo7a, and the baseline fluorescence intensity for each animal was calculated as the mean value of ROIs measured across 3 to 4 continuous single horizontal optical planes captured at the exact same cell body location. Individual data points in the right panel represent independent biological replicates (n = 5 per group). Data are plotted as mean ± S.D. Two-tailed Student’s *t*-test. ****P < 0.0001. Scale bar = 15 μm. **(B)** Representative whole-mount immunofluorescence surface images and corresponding semi-quantitative analysis of H3K36me2 modification levels in the sensory epithelium of adult *Nsd2^f/f^* and *Nsd2^Atoh1-KO^* mice. Confocal panels illustrate nuclear counterstaining via DAPI (blue), H3K36me2 enrichment (green), the canonical HC marker Myo7a (red), and the merged channels (Merge). Knockout mice display a severe reduction of downstream H3K36me2 levels within the HC rows. For cell-specific measurement, Regions of Interest (ROIs) were circumscribed around individual cell somas delineated by Myo7a, and the baseline value for each animal was calculated as the mean value of ROIs measured across 3 to 4 continuous single horizontal optical planes captured at the exact same cell body location. Individual data points represent independent biological replicates (n = 5 per group). Data are plotted as mean ± S.D. Two-tailed Student’s t-test, ***P < 0.001. Scale bar = 15 μm. **(C)** Auditory brainstem response (ABR) threshold measurements in adult *Nsd2^f/f^*and *Nsd2^Atoh1-KO^* mice across pure-tone frequencies ranging from 4 kHz to 32 kHz. Conditional knockout mice exhibit profound, generalized hearing impairment, characterized by significantly elevated ABR thresholds across all tested frequencies compared with control littermates. Individual data points plot the calculated threshold values from independent biological replicates (n = 6 per group). Data are presented as mean ± SEM. Two-way ANOVA with Tukey’s post hoc test, *P < 0.05, **P < 0.01, ***P < 0.001. **(D)** Representative ABR waveforms (waves I to V) from *Nsd2^f/f^* and *Nsd2^Atoh1-KO^* mice. Figure 2D presents the ABR recordings entirely in response to the click stimulus at a normalized suprathreshold level of 20 dB SPL above each individual animal’s auditory threshold. Each dashed line represents the raw trace recorded from a single individual animal (n = 6 per group), while the solid line represents the grand average waveform calculated across all animals within that specific genotype group to define peak latencies. Vertical scale bar = 2 μV. **(E–F)** Quantitative analysis of absolute latencies and interpeak latencies derived from click-evoked ABR waveforms at 20 dB SPL above the individual auditory threshold (*n* = 6 mice per group). Absolute latencies for waves I to V (E) and corresponding interpeak latencies for intervals I to II and II to III (F) were systematically extracted from the standardized suprathreshold state to eliminate the confounding effects of threshold shifts on waveform propagation. Pink lines denote control *Nsd2^f/f^* mice, and blue lines denote hair cell-specific knockout (*Nsd2^Atoh1-KO^*) mice. Data are presented as mean ± SEM. Two-way ANOVA with Tukey’s post hoc test, **P < 0.01, ***P < 0.001 **(G)** Distortion product otoacoustic emission (DPOAE) threshold profiles in adult *Nsd2^f/f^* and *Nsd2^Atoh1-KO^* mice. Quantitative assessment of 2f1-f2 DPOAE responses was performed across pure-tone frequencies from 4 kHz to 32 kHz. Conditional knockout mice show a profound deficit in outer hair cell electromotility, indicated by significantly elevated DPOAE thresholds across all tested frequencies compared with control littermates, except at 4 kHz. Individual data points plot the calculated threshold values from independent biological replicates (n = 6 per group). Data are presented as mean ±SEM. Two-way ANOVA with Tukey’s post hoc test, ***P < 0.001.

To evaluate whether NSD2 deficiency impairs auditory function, we recorded Auditory Brainstem Response (ABR) in adult *Nsd2^f/f^*and *Nsd2^Atoh1-KO^* mice (n=6 per group). As shown in Fig. 2C, *Nsd2^Atoh1-KO^* mice exhibited significantly elevated auditory thresholds across all tested frequencies from 4 kHz to 32 kHz compared to *Nsd2^f/f^* mice, with the most pronounced deficits observed in the 8 to 16 kHz range.

To characterize the physiological deficits underlying this hearing impairment, we analyzed suprathreshold ABR waveforms at the intensity of 20 dB SPL above the auditory threshold (Fig. 2D). The waveforms of *Nsd2^Atoh1-KO^* mice exhibited severely degraded reproducibility and markedly reduced amplitudes. Notably, this disruption was initiated at wave I, reflecting a severe deficit in peripheral auditory conduction and nerve synchrony that typically stems from compromised outer hair cell (OHC) electromotility and amplification (Fig. 2D). Concurrently, the peak latencies of subsequent waves underwent significant alterations and demonstrated a highly disorganized distribution across individual mutant mice, manifesting as a profound increase in inter-individual latency variability.

In addition, the wave latencies (I to V) were significantly prolonged in *Nsd2^Atoh1-KO^* mice compared to *Nsd2^f/f^*mice (Fig. 2E). Analysis of interpeak latencies further revealed a marked elongation of the I–II interval, whereas the II–III interval was decreased in the mutant group (Fig. 2F). Considering the prolonged latency of wave I alongside these altered interpeak intervals, there appeared early signs of a central auditory conduction disorder or other central auditory processing deficits ^36,37^. To identify the specific source of the hearing loss detected by ABR, we conducted distortion product otoacoustic emissions (DPOAE) testing (Fig.2G). The *Nsd2^Atoh1-KO^* mice exhibited significantly elevated DPOAE thresholds across the examined frequencies, with the most pronounced deficits observed at intermediate frequencies (8 kHz to 24 kHz) (Fig. 2G). These data together demonstrate that NSD2 deletion dampens outer hair cell electromotility, thereby disrupting peripheral auditory conduction and impairing overall hearing function.

### NSD2 deficiency leads to anatomical abnormality and an upregulation extracellular matrix based on single cell RNA sequencing

To investigate the anatomical alterations underlying the auditory deficits, we performed immunofluorescence staining for the HC marker Myo7a, coupled with DAPI and phalloidin labeling in adult cochlear tissues (Fig. 3A and Fig. S1D). The results revealed that while the total number and rows of HCs remained essentially unaltered (Fig. S2A), the anatomical organization of the organ of Corti was disrupted. Quantitatively, the span covering the three rows of OHCs was significantly widened in the *Nsd2^Atoh1-KO^* mice compared to *Nsd2^f/f^* mice (Fig. 3A). Notably, this spatial expansion was accompanied by a severe structural disorganization of the OHC nuclei. Instead of the canonical linear rows observed in controls, the OHC nuclei in the mutant cochlea displayed a disordered and misaligned arrangement, with the structural perturbation appearing most pronounced within the third row of OHCs (Fig. 3A). Concurrently, phalloidin staining revealed a visibly disorganized pattern of the hair bundles, characterized by irregular spacing between adjacent stereocilia clusters. Additionally, we observed an increase in the distance between two adjacent HC bundles (Fig. S2A). Together, these findings demonstrate that NSD2 deficiency perturbs the precise spatiotemporal alignment and cellular positioning of OHCs within the sensory epithelium without affecting initial HC survival.

**Fig. 3.**
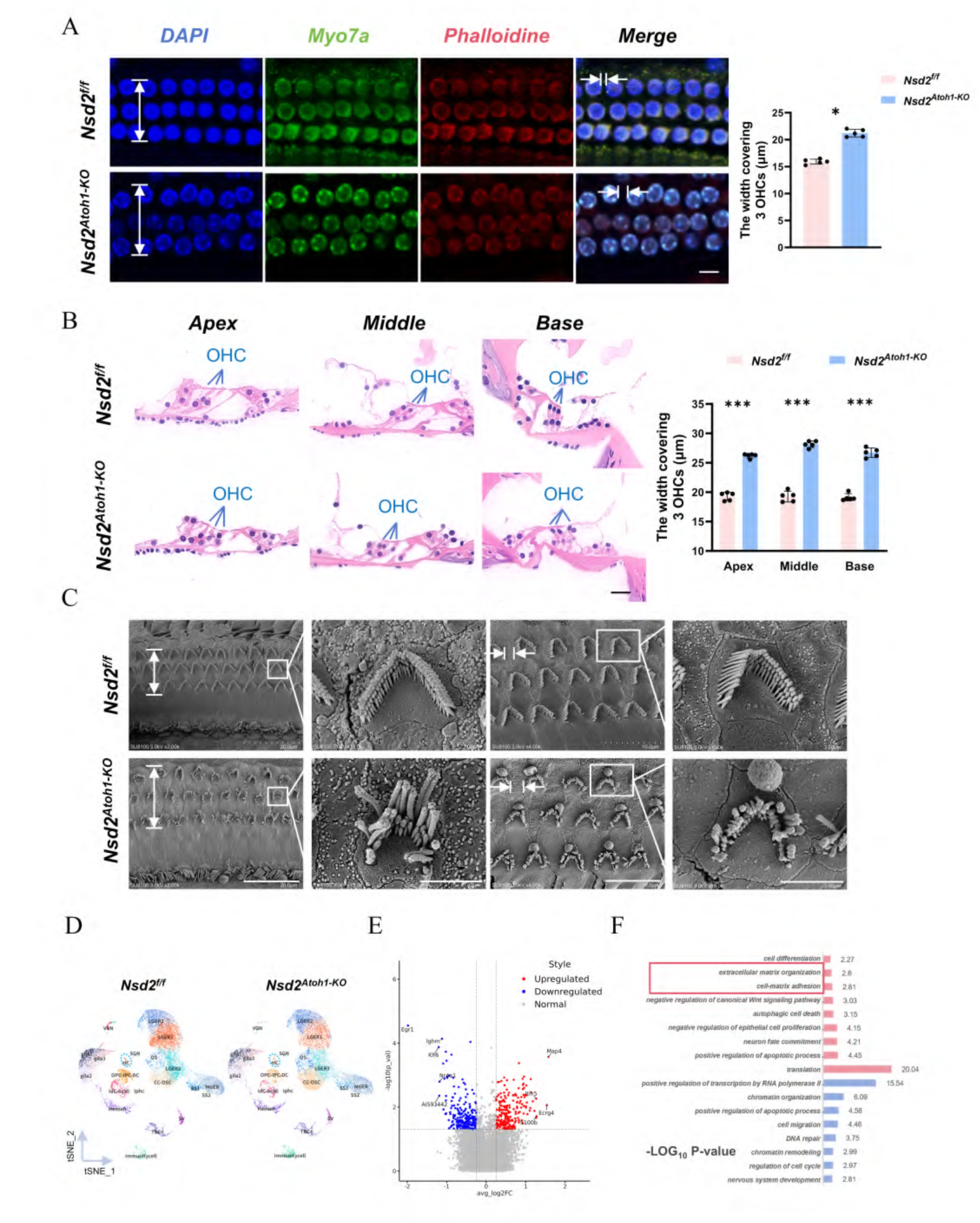
NSD2 deficiency leads to anatomical abnormality and an upregulation extracellular matrix based on single cell RNA sequencing. **(A)** Representative whole-mount immunofluorescence surface images and corresponding quantitative analysis of the sensory epithelium width in the cochlear middle turns of adult *Nsd2^f/f^* and *Nsd2^Atoh1-KO^* mice. Confocal panels illustrate nuclear counterstaining via DAPI (blue), Myo7a (green), phalloidin (red), and the merged channels (Merge). Vertical white double-headed arrows indicate the measurement strategy for determining the total width covering the 3 outer hair cell (OHC) rows quantified in the right panel. Horizontal white double-headed arrows in the Merge panels indicate the intercellular distance between adjacent OHCs, with its corresponding quantitative statistical analysis provided in supplementary figure S2A. Knockout mice exhibit a significant expansion and spacing disorganization of the OHC region. Individual data points in the right panel plot the calculated width values from independent biological replicates (n = 5 per group). Data are plotted as mean ± S.D. Two-tailed Student’s t-test, *P < 0.05. Scale bar = 10 μm. **(B)** Representative hematoxylin and eosin (H&E) stained cochlear cross-sections and corresponding quantitative analysis of the sensory epithelium width across different turns in adult *Nsd2^f/f^* and *Nsd2^Atoh1-KO^* mice. Histological panels illustrate the structural morphology of the Organ of Corti within the defined apical (Apex), middle (Middle), and basal (Base) turns. Bar graphs in the right panel represent the quantified width covering the 3 outer hair cell (OHC) rows across spatial turns. Individual data points plot the calculated measurement values from independent biological replicates (n = 5 per group). Data are plotted as mean ± S.D. Two-way ANOVA with Tukey’s post hoc test, ***P < 0.001. Scale bar = 25 μm.. **(C)** Representative scanning electron microscopy (SEM) micrographs illustrating the surface ultrastructure of stereocilia bundles within the cochlear middle turns of adult *Nsd2^f/f^*and *Nsd2^Atoh1-KO^* mice. Micrographs are organized sequentially across four columns from left to right. The first column presents low-magnification overviews of the sensory epithelium, where vertical white double-headed arrows define the total width covering the 3 outer hair cell (OHC) rows. The third column displays medium-magnification fields, where horizontal white double-headed arrows indicate the precise inter-bundle distance between adjacent OHC stereocilia bundles quantified in supplementary figure S2B. The second and fourth columns show high-magnification close-up views of the individual hair bundles corresponding to the highlighted white boxed regions, demonstrating severe stereocilia disorganization, bundle fusion, and structural degeneration in the conditional knockout mice. Scale bars = 20 μm (first column), 3 μm (second column), 10 μm (third column), and 3 μm (fourth column). **(D)** Two-dimensional t-distributed stochastic neighbor embedding (t-SNE) plots demonstrating the single-cell transcriptional landscape of the P7 mouse cochlea in *Nsd2^f/f^* and *Nsd2^Atoh1-KO^*groups via scRNA-seq analysis. Individual dots represent distinct single-cell transcriptomes color-coded by their corresponding cell-type identities. The sensory hair cell (HC) cluster is explicitly highlighted and delineated by blue dashed circles based on the expression profiles of canonical hair cell marker genes. **(E)** Volcano plot displaying differentially expressed genes (DEGs) identified in the hair cell (HC) cluster between *Nsd2^f/f^* and *Nsd2^Atoh1-KO^* mice. The horizontal axis indicates the log2 fold change (avg_log2FC) of gene expression, and the vertical axis indicates the statistical significance (-log10 P-value). Red dots represent significantly upregulated genes, blue dots represent significantly downregulated genes, and grey dots indicate genes with no significant transcriptomic alterations (Normal). **(F)** Gene Ontology (GO) enrichment analysis of differentially expressed genes (DEGs) within the hair cell (HC) cluster of conditional knockout mice compared with control littermates. Pink horizontal bars represent significantly enriched biological process terms for upregulated DEGs, while blue horizontal bars represent enriched terms for downregulated DEGs. The horizontal scale explicitly reflects the statistical significance scores denoted as -LOG_10_ P-value, with numerical values provided adjacent to each bar. Red boxes highlight the preferential enrichment of cell-matrix adhesion and extracellular matrix organization.

To evaluate the cross-sectional pathological alterations and potential cellular rearrangement suggested by the architectural disorganization, we examined hematoxylin and eosin (H&E) stained cochlear sections across the apical, middle, and basal turns (Fig. 3B). Morphometric assessment of H&E-stained sections demonstrated an increased width spanning the three rows of OHCs in Nsd2Atoh1-KO cochleae (Fig. 3B). Scanning electron microscopy (SEM) images revealed that the stereocilia bundles in *Nsd2^Atoh1-KO^* mice exhibited a distinctly flattened shape compared to those in control mice (Fig. 3C). The arrangement of OHC stereocilia was somewhat disorganized in *Nsd2^Atoh1-KO^* mice compared with *Nsd2^f/f^*mice, and the W-shaped stereocilia bundles in outer hair cells exhibited a distinct morphological change. Most OHC in *Nsd2^Atoh1-KO^* mice displayed less distinct and rounded stereocilia bundles, with typical W-shaped bundles being present in very few OHC of *Nsd2^Atoh1-KO^* mice, which indicated that the cell-intrinsic polarity was deficient in *Nsd2^Atoh1-KO^* hair cells. We also counted the width of the three rows of OHC, the number of hair cells and the spacing between the stereocilia bundles (Fig. S2A).

Quantitative assessment of these ultrastructural features confirmed that while the total number of HCs in the same given width remained unchanged, the row span covering the three rows of OHCs was significantly expanded (Fig. 3C). Furthermore, statistical measurement of the inter-bundle distance verified an increased physical distance between adjacent OHC hair bundles (Fig. S2B). This localized spacing alteration directly corroborates the expanded interstitial spaces observed in H&E cross-sections (Fig. 3B), collectively reflecting an overall architectural misalignment within the sensory rows rather than the addition of supernumerary cell types.

To evaluate the molecular alterations parallel to these structural deficits, we performed RT-qPCR to examine the transcription of key genetic components governing HC maturation and auditory function in P4 mice (Fig. S2C). The results demonstrated a significant reduction in the expression levels of several critical transcripts, including *Myo3a*, *Tmc1*, *Espnl*, *Tomt*, and *Gjb2*, within the P7 *Nsd2^Atoh1-KO^* cochlea compared to controls (Fig. S2C).

Given that loss or variation in each of these genes independently causes deafness, their combined downregulation, together with these morphological abnormalities, likely underlies the observed hearing impairment.

To investigate the possible transcriptome changes in HCs, we performed single cell RNA sequencing (scRNA-seq) on freshly dissected P7 murine cochlear tissues (Fig. 3D). Unsupervised clustering and t-SNE visualization resolved distinct cellular lineages, with the HC cluster precisely validated by the enriched expression of canonical markers including *Myo7a* and *Atoh1* (Fig. 3D and Fig. S2D). Differential expression analysis within the HC cluster identified a total of 438 differentially expressed genes (DEGs) post-NSD2 ablation, with 216 genes upregulated and 222 genes downregulated (Fig. 3E). Gene Ontology (GO) enrichment analysis of scRNA-seq revealed that the downregulated DEGs were predominantly involved in chromatin organization, regulation of cell cycle, and nervous system development (Fig. 3F). Conversely, upregulated DEGs were significantly enriched in pathways associated with cell differentiation, cell-matrix adhesion, and autophagic cell death (Fig. 3F and Fig. S2E). Among the altered pathways, it is particularly noteworthy that the extracellular matrix (ECM)-related pathways were significantly upregulated in the mutant HCs (Fig. 3F and Fig. S2E). To validate the aberrant activation of the ECM cascade, we evaluated the expression profiles of key ECM-related transcripts within the cochleae via RT-qPCR (Fig. S2E). As shown in Fig. S2D, the transcription of *Col4a4*, *Egfl6*, *Mmp15*, and *Npnt* was significantly elevated in the mutant group, closely replicating the trends observed in the scRNA-seq datasets. This excessive accumulation of ECM components directly mirrors our histological observations of the expanded width of the OHC region and increased interstitial voids (Fig. 3A and B). The excessive synthesis of ECM proteins likely drives the expansion of interstitial spaces, suggesting a functional link to auditory dysfunction. Taken together, these findings suggest that the loss of NSD2 triggers an anomalous remodeling of the extracellular matrix, potentially disrupting the homeostatic biomechanical properties of the cochlear microenvironment.

### NSD2-deficiency promotes expression of key ECM protein collagen in cochlear hair cells

To provide direct evidence that ECM production can be regulated by NSD2, we decided to establish in vitro model by knocking out the *Nsd2* gene in HC-like HEI-OC1 cells. Using lentiviral vectors encoding two independent sgRNA sequences targeting *Nsd2*, we established stable *Nsd2*-knockout HEI-OC1 cells (*Nsd2 Sg*), alongside empty lentiCRISPR v2 vector-transduced cells as controls (*Cv2*). Immunofluorescence revealed a marked decrease in NSD2 and H3K36me2 expression (Fig. S3 A-D), confirming the successful establishment of the NSD2-partial knockout HEI-OC1 (Fig. S3A-D). As shown in Fig. 4A, the loss of NSD2 led to a significant decrease in attachment area of cells. Furthermore, RT-qPCR results showed that the expression of HC development genes, including *Myo3a*, *Tmc1*, *Nsd2*, *Notch1*, *Espnl*, and *Tomt*, was significantly downregulated in *Nsd2 sg* cells (Fig. S3E).

**Fig. 4.**
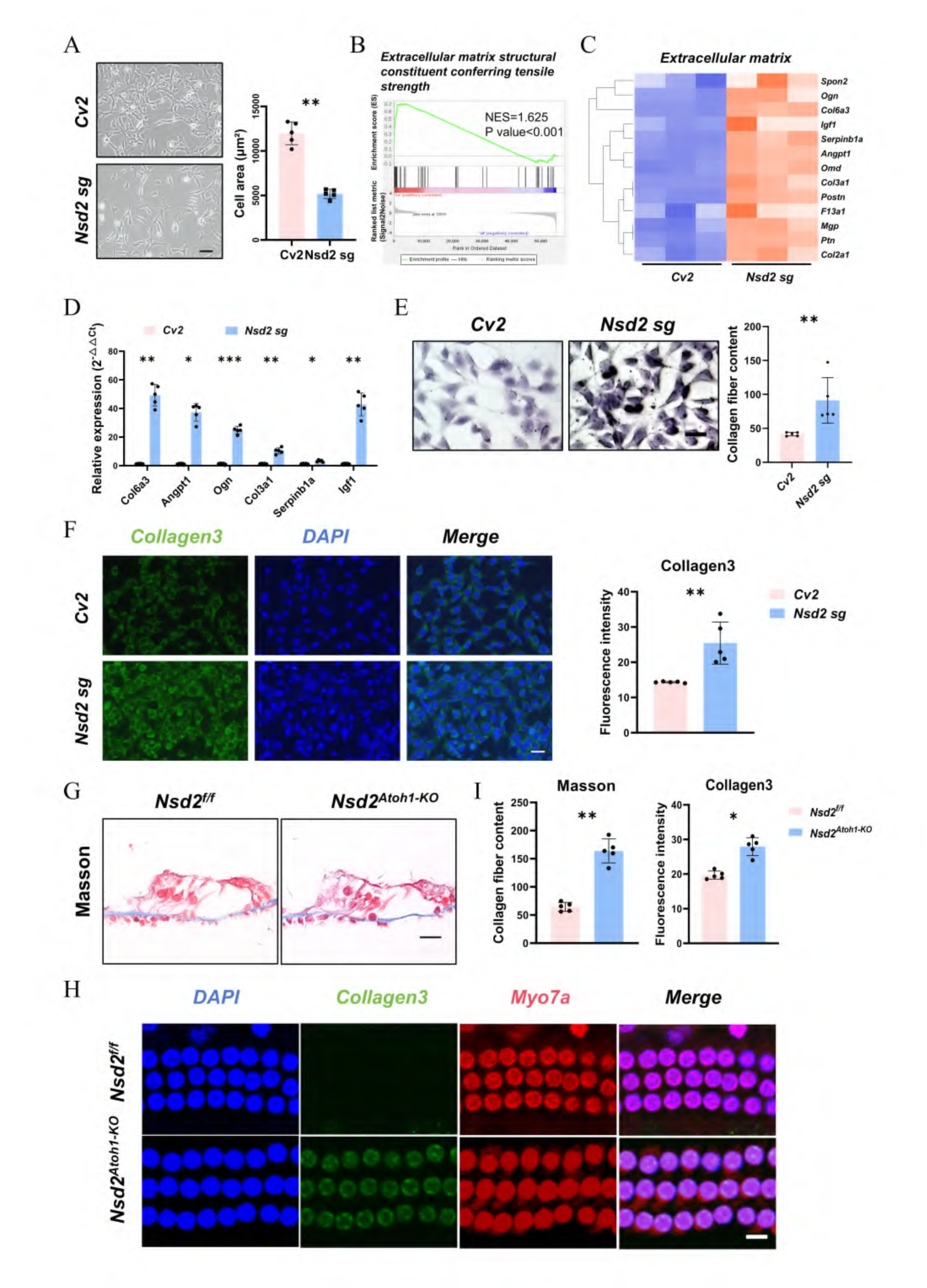
NSD2-deficiency promotes expression of key ECM protein collagen in cochlear hair cells. **(A)** Representative phase-contrast images and corresponding quantitative analysis of cell surface area in control (*Cv2*) and CRISPR-mediated Nsd2 knockout (*Nsd2 sg*) cells. Phase-contrast micrographs in the left panels illustrate the distinct cellular morphology and spreading patterns of each experimental group. The bar graph in the right panel represents the quantified cross-sectional cell area expressed in μm^2^. Individual data points plot the calculated cell area values obtained from independent biological replicates (n = 5 per group). Data are presented as mean ± S.D. Two-tailed Student’s t-test, **P < 0.01. Scale bar = 50 μm. **(B)** Gene set enrichment analysis (GSEA) plot demonstrating the coordinated upregulation of extracellular matrix (ECM) related pathways in CRISPR-mediated *Nsd2 sg* cells compared with control cells. The analysis reveals a significant enrichment of differentially expressed transcripts within the specific gene set designated as extracellular matrix structural constituent conferring tensile strength. The green line traces the calculated running enrichment score (ES) across the ranked list of genes. Statistical parameters indicate highly significant transcriptional enrichment profile with a normalized enrichment score (NES = 1.625) and statistical significance (P value < 0.001, false discovery rate FDR < 0.05). **(C)** Hierarchical clustering heatmap summarizing the relative expression profiles of selected ECM related genes in *Cv2* and *Nsd2 sg2* cells (n = 3 independent biological replicates per group). Rows represent individual ECM-associated transcripts including canonical structural components such as *Col4a1, Fn1, Lama4,* and *Lamb1*, while columns represent independent sample replicates. The color scale bar at the upper right indicates relative gene expression levels based on Row Z-score normalization, where red denotes high expression and blue denotes low expression. Knockout of *Nsd2* triggers a robust and synchronized transcriptional upregulation of key ECM structural constituents. **(D)** RT-qPCR analysis validating the relative mRNA expression levels of candidate ECM related genes in *Cv2* and *Nsd2 sg2* cells. Examined transcripts include *Col6a3, Angpt1, Ogn, Col3a1, Serpinb1a*, and *Igf1*. Transcript levels are calculated utilizing the comparative 2^-ΔΔCT^ method and normalized to internal control thresholds. Individual data points plot the calculated relative fold-change values from independent biological replicates (n = 5 per group). Data are presented as mean ± S.D. Two-tailed Student’s t-test, *P < 0.05, **P < 0.01,***P < 0.001. **(E)** Representative Masson’s trichrome staining and corresponding quantitative analysis of collagen fiber content in *Cv2* and *Nsd2 sg2* cells. Micrographs in the left panels illustrate the intracellular and extracellular deposition of collagen fibers. The bar graph in the right panel represents the quantified relative collagen fiber content reflecting histological ECM accumulation. Individual data points plot the calculated intensity values obtained from independent biological replicates (n = 5 per group). Data are presented as mean ± S.D. Two-tailed Student’s t-test, **P < 0.01. Scale bar = 10 μm. **(F)** Representative immunofluorescence images and corresponding quantitative analysis of Collagen3 expression in *Cv2* and *Nsd2 sg2* cells. Micrographs display specific immunostaining for Collagen3 (green) and counterstaining for cell nuclei via DAPI (blue), with merged profiles shown in the rightmost columns. The bar graph plots the quantified relative fluorescence intensity reflecting structural ECM alterations. Individual data points indicate values obtained from independent biological replicates (n = 5 per group). Data are presented as mean ± S.D. Two-tailed Student’s t-test, **P < 0.01. Scale bar = 25 μm. **(G)** Representative Masson’s trichrome staining of the Organ of Corti sections from adult *Nsd2^f/f^* and *Nsd2^Atoh1-KO^*mice. Histological micrographs demonstrate the structural distribution and deposition of collagen fibers within the basilar membrane beneath the sensory epithelium. Images are representative of n = 5 per experimental group. Scale bar = 25 μm. **(H)** Representative whole-mount immunofluorescence images of Collagen3 expression within the cochlear basilar membrane and sensory epithelium of *Nsd2^f/f^*and *Nsd2^Atoh1-KO^* mice. Confocal micrographs display specific staining for Collagen3 (green), Myo7a (red) to delineate hair cell boundaries, and nuclei via DAPI (blue). Images are representative of n = 5 per group. Scale bar = 25 μm. **(I)** Quantitative analysis of the relative collagen fiber content from Masson’s trichrome staining in (G) and the relative immunofluorescence intensity of Collagen3 in (H) comparing *Nsd2^f/f^* and *Nsd2^Atoh1-KO^* cochleae. Individual data points plot the quantified values calculated from independent biological replicates (n = 5 per experimental group). Data are presented as mean ± S.D. Two-tailed Student’s t-test, *P < 0.05, **P < 0.01.

To investigate whether NSD2 deficiency leads to ECM accumulation in HC-like cells, we conducted RNA-seq on *Cv2* and *Nsd2 sg* HEI-OC1. Gene Set Enrichment Analysis (GSEA) and heatmap showed the upregulation of the ECM pathway following NSD2 loss (Fig. 4 B and C). RT-qPCR validation confirmed the elevated transcription of ECM-related components in the cell line dataset, which consisted of *Collagen3, Angpt1, Ogn, Col6a3, Serpinb1a, Igf1, Bcam, Npnt, Tecta, Ccdc80,* and *Col4a4*. (Fig. 4D). Similarly, the upregulation of scRNA-seq targets, including Bcam, Npnt, Tecta, Ccdc80, and Col4a4, was also reproduced in *Nsd2 sg* cells (Fig. S3F).

Considering that collagen is a major component of the ECM in the cochlea, subsequently, we focused more on collagen that serves as a representation of ECM. Masson’s trichrome staining demonstrated a significant increase in total collagen fiber content within *Nsd2 sg* cells compared to controls (Fig. 4E). To pinpoint the specific collagen types driving this matrix accumulation, we next investigated Collagen 3. While Collagen 3 is minimally expressed in the cochlea under physiological conditions, our sequencing data highlighted it as a prominent pathological indicator of NSD2 deficiency. Immunofluorescence further verified that NSD2 loss prominently induced the accumulation of Collagen 3 in HEI-OC1 cells (Fig. 4F).

To determine if this pathological cascade operates *in vivo*, we performed parallel histological evaluations on cochlear sections from *Nsd2^f/f^* and *Nsd2^Atoh1-KO^*mice (Fig. 4 G and I). Consistent with the *in vitro* findings, Masson’s staining revealed robust ECM accumulation in mutant cochlear HCs (Fig. 4G and I), accompanied by a profound increase in collagen 3 immunoreactivity within the sensory epithelium (Fig. 4H and I). Strikingly, this intracellular collagen deposition was not restricted to collagen 3. Supplementary immunofluorescence confirmed a marked accumulation of other basement membrane and interstitial components (including collagen 1, 4, and 5) within the mutant hair cells (Fig. S4A). Collectively, these in vitro and in vivo data demonstrate that NSD2 deficiency drives an aberrant overproduction and accumulation of key ECM collagens within cochlear hair cells.

### ECM intervention mitigates NSD2 deficiency-mediated hearing loss and alters cellular stiffness-dependent cellular behavior

To explore the therapeutic potential of targeting deregulated ECM deposition in vivo, we first implemented AAV-mediated RNA interference to achieve targeted knockdown of Ogn or Col6a3 within the cochlear HCs of 2-week-old *Nsd2^Atoh1-KO^* C57BL/6J mice (validated in Fig. S5A). ABR measurements revealed that silencing either *Ogn* or *Col6a3* significantly and partially restored hearing thresholds across multiple frequencies in mutant mice (Fig. 5A). Immunofluorescence staining for Myo7a and subsequent SEM demonstrated that this ECM intervention effectively mitigated the structural disorganization of the sensory epithelium. This structural rescue was characterized by a significant reduction in the width covering the 3 OHC rows compared to the *sh-NC* mutant controls (Fig. 5B, 5C, and Fig. S5B), approaching the physiological arrangement observed in *Nsd2^f/f^* controls. Immunofluorescence analysis further confirmed a prominent reduction in aberrant collagen 3 accumulation following *Ogn* or *Col6a3* knockdown, largely reversing the matrix accumulation toward physiological controls (Fig. 5B). Crucially, restoring NSD2 expression via AAV-mediated gene therapy (validated in Fig. S5D) similarly reduced pathological Collagen 3 deposition, rescued the OHC arrangement anomalies, and ameliorated hearing loss in mutant mice (Fig. 5D-F and Fig. S5E). These *in vivo* findings demonstrate that targeted ECM intervention or genetic restoration can effectively rescue hearing damage caused by NSD2 deficiency.

**Fig. 5.**
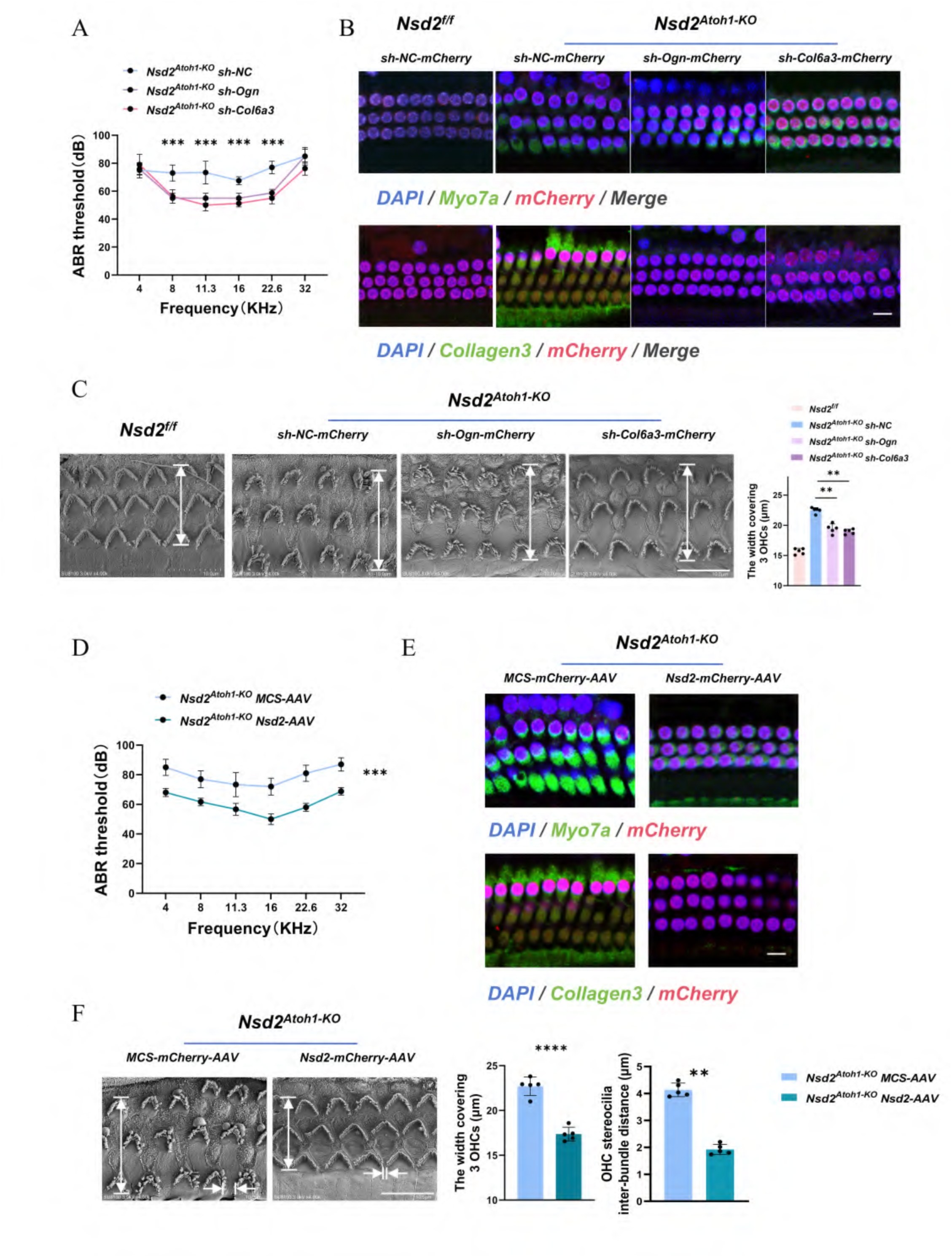
ECM intervention mitigates NSD2 deficiency-mediated hearing loss and alters cellular stiffness-dependent cellular behavior. **(A)** ABR threshold curves across specified frequencies ranging from 4 to 32 KHz comparing control and gene-targeted rescue cohorts. Hair cell-specific knockout mice were treated with adeno-associated viruses delivering non-target control sequences or specific hairpins, designating the groups as *Nsd2^Atoh1-KO^ sh-NC*, *Nsd2^Atoh1-KO^ sh-Ogn*, and *Nsd2^Atoh1-KO^ sh-Col6a3*. Viral knockdown of either upregulated ECM factor significantly reduces auditory thresholds and rescues functional hearing capacity under NSD2 deficiency conditions. Data are presented as means ± SEM (n = 3 per experimental group). Two-way ANOVA followed by Bonferroni’s post hoc test, ***P < 0.001 versus the *sh-NC* group. **(B)** Representative whole-mount immunofluorescence confocal images of the organ of Corti demonstrating structural preservation and ECM remodeling. Upper rows illustrate specific staining for the canonical hair cell marker Myo7a (green), cell nuclei via DAPI (blue), and successful viral transduction via mCherry (red). Lower rows display specific immunolabeling for Collagen3 (green) to assess ECM deposition across control (*Nsd2^f/f^*) and knockout (*Nsd2^Atoh1-KO^*) groups under indicated treatment conditions. Scale bar = 25 μm. **(C)** Representative scanning electron microscopy (SEM) micrographs and corresponding quantitative analysis measuring the structural width covering three rows of outer hair cells (OHCs) to evaluate spatial organization. Cochleae from adult control (*Nsd2^f/f^*) mice are compared with hair cell-specific knockout cohorts treated with control or targeted viral vectors, designated as *Nsd2^Atoh1-KO^ sh-NC*, *Nsd2^Atoh1-KO^ sh-Ogn,* and *Nsd2^Atoh1-KO^ sh-Col6a3*. White double-headed vertical arrows mark the measured radial width spanning across the three OHC rows. Knockdown of either upregulated ECM constituent effectively ameliorates the abnormal epithelial spreading and broadening defects observed under NSD2 deficiency. Individual data points plot the calculated width values in μm from independent biological replicates (n = 5 per experimental group). Data are presented as mean ± S.D. One-way ANOVA followed by Tukey’s post hoc test, **P < 0.01, ***P < 0.001. Scale bar = 10 μm. **(D)** ABR threshold curves across specified frequencies ranging from 4 to 32 KHz evaluating the functional rescue of hearing loss via exotic gene supplement. Hair cell-specific knockout mice were delivered with control or Nsd2-reexpressing viral vectors, designating the cohorts as *Nsd2^Atoh1-KO^ MCS-AAV* and *Nsd2^Atoh1-KO^ Nsd2-AAV*. Reexpression of *Nsd2* significantly reduces auditory thresholds and rescues the hearing impairment phenotype caused by endogenous NSD2 deficiency. Data are presented as mean ± S.D. Two-way ANOVA followed by Bonferroni’s post hoc test, ***P < 0.001. **(E)** Representative whole-mount immunofluorescence confocal micrographs of the organ of Corti demonstrating cellular preservation and ECM containment after gene rescue. Upper rows show specific immunostaining for Myo7a (green), nuclei via DAPI (blue), and viral transduction markers via mCherry (red). Lower rows show specific immunolabeling for Collagen3 (green) to monitor ECM deposition within the sensory epithelium under indicated virus delivery. Scale bar = 25 μm. **(F)** Representative SEM micrographs and corresponding quantitative analyses detailing the morphological preservation of outer hair cell (OHC) microarchitecture. White double-headed vertical arrows in the left panels delineate the total radial width covering three rows of OHCs, which is quantified in the left bar graph. Horizontal opposing white arrowheads specify the gaps separating individual stereocilia bundles, quantified as the OHC stereocilia inter-bundle distance in the right bar graph. Virally mediated *Nsd2* re-expression (*Nsd2^Atoh1-KO^ Nsd2-AAV*) effectively counteracts the aberrant epithelial spreading and restabilizes both the cellular row width and inter-bundle spatial intervals back to tightly packed physiological parameters compared with empty vector controls (*Nsd2^Atoh1-KO^ MCS-AAV).* Individual data points plot the calculated micrometric values obtained from independent biological replicates (n = 5 per experimental group). Data are presented as mean±S.D. Two-tailed Student’s t-test, **P < 0.01, ****P < 0.0001. Scale bar = 10 μm.

To dissect the cellular and mechanical mechanisms underlying this matrix-mediated damage, we varied substrate stiffness to evaluate how cells respond to different rigidity microenvironments. *Cv2* and *Nsd2 sg* HEI-OC1 cell areas were evaluated on Matrigel substrates with varied stiffnesses (1.25 kPa, 0.97 kPa, 0.65 kPa, and 0.42 kPa) (Fig. S6A). To find out the physiological stiffness of native cochlear tissue, we performed atomic force microscopy (AFM) analysis of normal cochlear hair cells, which showed a range approximately from 1.05 to 1.25 kPa. Although both groups achieved stable attachment within 24 hours, *Cv2* cells displayed significantly enhanced cell area compared to *Nsd2 sg* cells on physiological stiffness (1.25 kPa and 0.97 kPa) (Fig. S6 A-C). Conversely, on softer matrices (0.65 kPa and 0.42 kPa), *Cv2* cells exhibited a restricted cell area, whereas *Nsd2 sg* cells demonstrated significantly increased cell areas (Fig. S6 A-C). To further evaluate whether these matrix stiffness alterations impact cell motility, we performed Transwell migration assays under different stiffness conditions. The analysis revealed that NSD2-deficient HEI-OC1 cells displayed altered migratory behaviors and cellular dispersion patterns compared to the control groups (Fig. S6D). To further corroborate this stiffness-dependent behavior, we evaluated NSD2-overexpressing HEI-OC1 cells (*Nsd2 oe*) under normal (0.97 kPa) and low (0.42 kPa) stiffness conditions. *Nsd2 oe* cells showed an increased cell area at 0.97 kPa but displayed a severely diminished cell area at 0.42 kPa (Fig. S7A).

RT-qPCR analysis confirmed that *Nsd2* deletion sustained elevated expression of *Col4a4*, *Ogn*, and *Col3a1*, whereas *Nsd2* overexpression significantly suppressed these ECM components across different stiffness levels (Fig. S7B). Finally, to determine whether the NSD2-deficient phenotype directly stems from this deregulated ECM deposition, we genetically ablated *Ogn* or *Col6a3* in *Nsd2 sg* cells. Dual knockout of *Nsd2* with either matrix gene rescued the cell area on physiological stiffness substrates (Fig. S8 A and B; knockout efficiencies validated in Fig. S8C), and restored the expression of essential cochlear functional markers (Fig. S8D). Collectively, these data confirm that NSD2 deficiency impairs how cells respond to different rigidity microenvironments and suppresses cell migration, which is associated with matrix overproduction and can be effectively mitigated by targeted ECM intervention.

### Loss of H3K36me2 causes a genome-wide alteration in chromatin accessibility that correlates with increased extracellular matrix transcription

To dissect the downstream molecular mechanisms altered by NSD2 deficiency in auditory cells, we performed transcriptomic and epigenetic profiling using HC-like HEI-OC1 cells ^38,39^. The RNA sequencing data revealed 443 differentially expressed genes (DEGs), with 118 genes upregulated and 325 genes downregulated following NSD2 knockdown (Fig. S9 A and B). GO and GSEA pathway analysis revealed that showed that these DEGs were primarily enriched in cell differentiation, neural apoptosis, negative regulation of cell development, and gene expression processes (Fig. 6A, S9C-E).

**Fig. 6.**
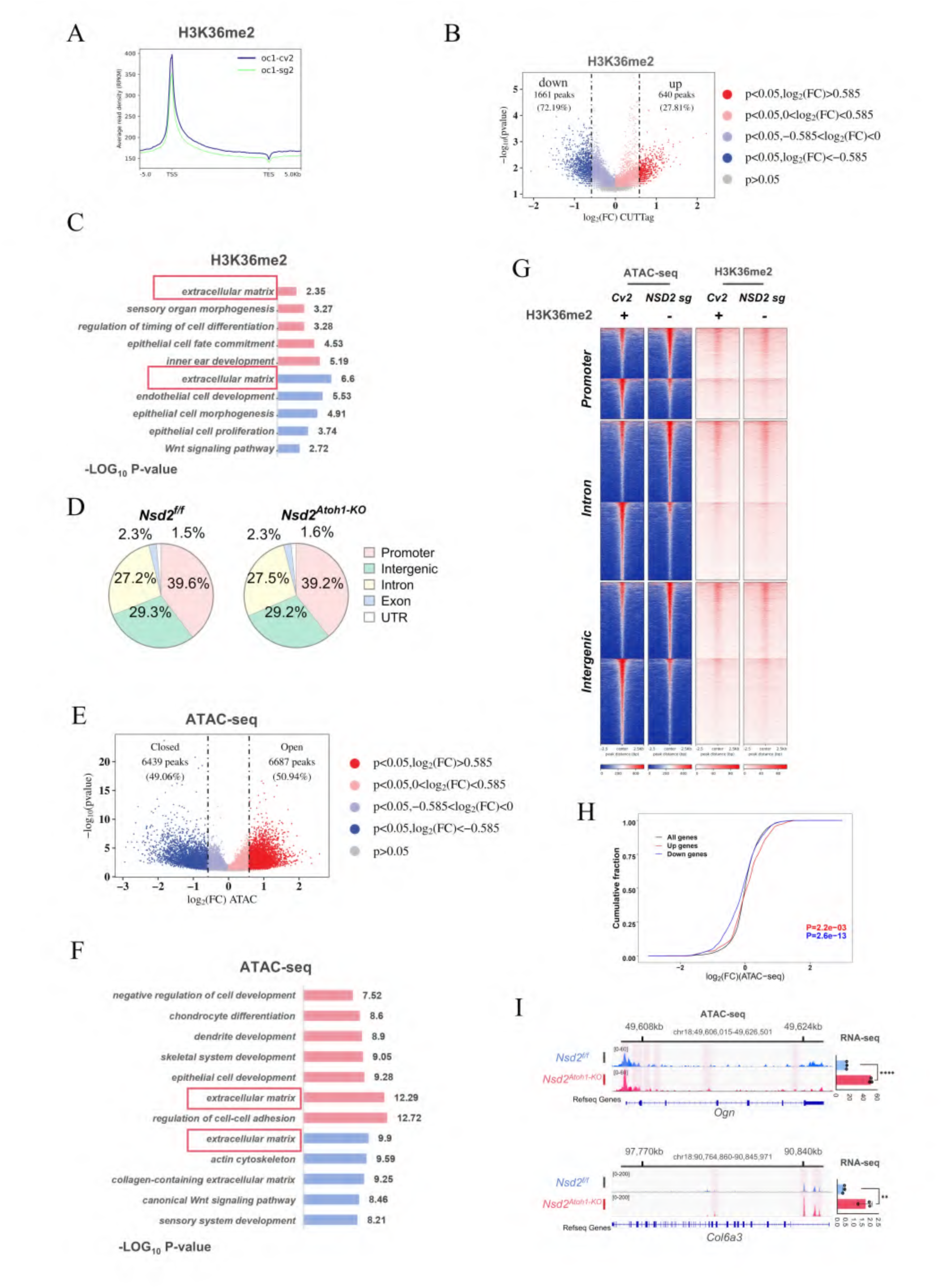
Loss of H3K36me2 causes a genome-wide alteration in chromatin accessibility that correlates with increased extracellular matrix transcription. **(A)** Metaplots showing the normalized average levels of H3K36me2 across gene bodies comparing control to *Nsd2 sg* cells by CUT&Tag; TSS, transcription start site; TES, transcription end site. **(B)** Volcano plots showing changes in CUT&Tag comparing *Cv2* and *Nsd2 sg* cells. Peaks with differential enrichment for H3K36me2 (FDR < 0.05) are highlighted. Loci exhibiting increased H3K36me2 occupancy were annotated as transcriptionally activated domains, while those with decreased occupancy were designated as repressed loci. The number of peaks with significant changes (FDR < 0.05 and log2 (FC) > 0.585) in H3K36me2 is shown. **(C)** GO enrichment analysis of differential loci identified via H3K36me2 CUT&Tag sequencing to investigate epigenetic regulation mechanisms. The bar graph plots the enriched biological processes and structural categories ranked by their statistical significance expressed as -LOG_10_ P-value values. Red rectangular boxes highlight the significant enrichment of loci associated with the ECM category. Additional highly enriched functional groups closely align with developmental programs including inner ear development, sensory organ morphogenesis, and epithelial cell fate commitment, directly coupling NSD2-mediated histone modifications to auditory gene regulatory networks. **(D)** Pie charts show the percentage of differentially accessible ATAC-seq peaks (FDR < 0.05) at promoter, intronic, intergenic, exonic, and UTR regions. **(E)** Volcano plot of ATAC-seq peaks comparing *Cv2* and *Nsd2 sg* cells. The chromatin states are relative to wild-type baselines. “Open chromatin” (red peaks) refers to the genomic regions exhibiting significantly increased accessibility in mutant cells and “closed chromatin” (blue peaks) represents the regions with reduced accessibility in mutant cells. The number of peaks with significant changes (FDR < 0.05 and log2 (FC) > 0.585; n = 12,136 peaks) is shown. **(F)** GO enrichment analysis of differentially accessible regions (DARs) identified via ATAC-seq to determine global chromatin accessibility alterations. The bar graph illustrates significantly enriched biological processes and cellular components ranked by their statistical significance expressed as -LOG_10_ P-value. Red rectangular boxes emphasize the highly significant enrichment of genomic loci residing within the ECM pathway. Functional clusters are prominently mapped to core structural and developmental programs including regulation of cell-cell adhesion, collagen-containing ECM, and sensory system development, indicating that NSD2 depletion triggers massive chromatin remodeling that directly licenses the transcriptional activation of ECM structural constituents. **(G)** Heat maps of differentially accessible ATAC-seq peaks (FDR < 0.05 and log2(FC) > 0.585; n = 12,136) grouped by localization at promoter, intron, and intergenic regions and CUT&Tag signals for the indicated histone modifications in the same regions of ATAC-seq peaks.Go term analysis of ATAC-seq. **(H)** Distribution of chromatin accessibility changes associated with significantly upregulated (red) or downregulated (blue) genes. P values were calculated using a one-sided Kolmogorov–Smirnov (KS) test comparing peaks associated with differentially expressed genes to all genes **(I)** Left, IGV tracks of open chromatin at *Ogn* or *Col6a3* (with y-axis scales as indicated). Right, normalized bulk RNA-seq counts of *Ogn* or *Col6a3* in *Cv2* and *Nsd2 sg* cells (n = 3).

**Fig. 7.**
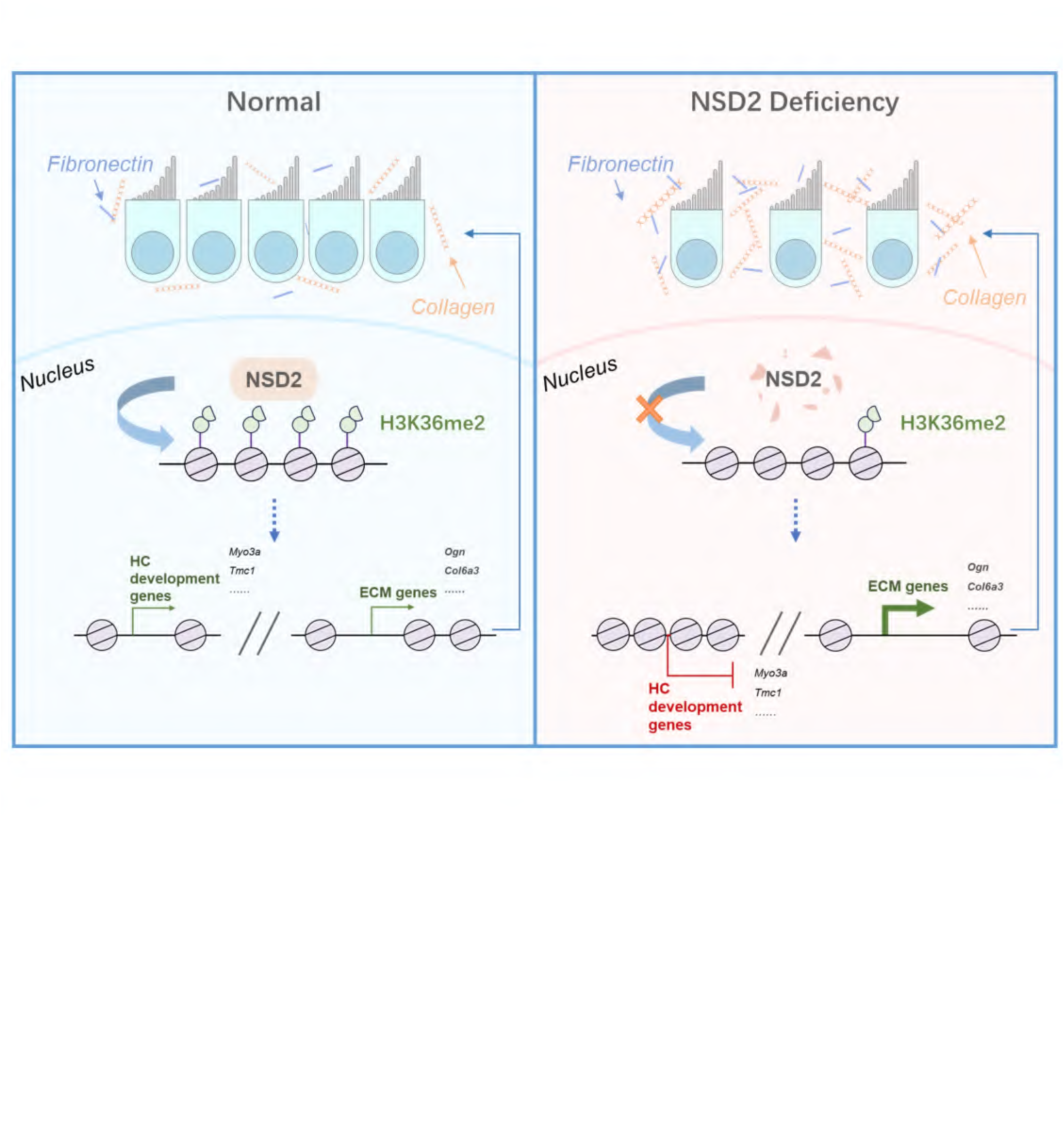
NSD2 deficiency disrupts the epigenetic landscape of cochlear HCs to enhance ECM expression and perturb cochlear HC development.

To directly investigate the primary epigenetic alterations responsible for these transcriptomic shifts, we performed CUT&Tag profiling using H3K36me2-specific antibodies. NSD2 loss induced a global, genome-wide decrease in H3K36me2 deposition spanning both promoter and gene body regions (Fig. 6A), which is consistent with NSD2 knockout results in other organs^40–42^. Differential analysis revealed that the loss of H3K36me2 led to 72.19% downregulated genes (Fig. 6B), with these down-regulated target genes enriched in developmental pathways such as sensory organ morphogenesis, inner ear development, and the Wnt signaling pathway (Fig. 6C). While 27.81% genes were upregulated, including those associated with epithelial cell fate and inner ear development pathways. (Fig. 6 B and C).

However, because the loss of NSD2-mediated H3K36me2 can extensively influence the chromatin landscape and potentially cross-talk with multiple overlapping histone modifications or remodeling complexes, tracking this single histone mark alone is insufficient to fully resolve the broader structural state of these loci. To better understand the downstream molecular mechanism and directly assess global chromatin accessibility, we subsequently performed ATAC sequencing. The ATAC sequencing data revealed that NSD2 loss resulted in altered chromatin accessibility (Fig. 6 D and E), extensively reshaping the global landscape. The alterations in chromatin accessibility caused by NSD2 deficiency were directly correlated with the upregulation of negative regulators of cell development and extracellular matrix pathways, as well as the downregulation of sensory system development pathways (Fig. 6F). In addition, we observed global alterations in chromatin openness within specific genomic regions, including the promoter, intron, and intergenic regions, which correlated with the local H3K36me2 distribution pattern (Fig. 6G).

To determine how these structural alterations dictate transcriptional output, we integrated the ATAC-seq and RNA-seq datasets. A positive correlation observed between the chromatin accessibility changes chromatin accessibility changes and transcriptional output induced by *Nsd2* loss (Fig. 6H); specifically, upregulated genes exhibited a more open chromatin, whereas downregulated genes overlapped with regions of reduced accessibility (Fig. 6H).

To identify the specific downstream networks driving HC dysfunction, we performed transcription factor (TF) motif analysis within these differentially accessible ATAC-seq peaks using FIMO29 (Fig. S10A). Among the top enriched TFs, those associated with open chromatin regions in NSD2-deficient cells were primarily linked to developmental arrest, whereas TFs enriched in closed regions included critical embryonic and neuronal development regulators, such as members of the HOX family, FOXL2, and PAX2 (Fig. S10A).

It is noteworthy that the extracellular matrix pathway was also enriched across our RNA-seq, CUT & Tag, and ATAC detection (Fig. 6C, F and Fig. S9B), confirming that aberrant matrix deposition is deeply integrated with the epigenetic and structural remodeling process. And chromatin accessibility changes at loci encoding ECM, such as *Ogn* and *Col6a3*, directly correlate with their transcriptional upregulation, demonstrating a locus-specific alignment between chromatin openness and transcriptional output (Fig. 6I). A similar correlation was observed for other essential matrix components, including *Col3a1* and *Ccdc80* (Fig. S10B).

Collectively, these multi-omic findings reveal that the loss of NSD2 decreases H3K36me2, leads to a global alteration in chromatin accessibility. This structural remodeling enhances the extracellular matrix transcriptional output in cochlear HCs, which in turn affects the hearing function.

## Discussion

Auditory impairment is one of the most prevalent disorders of the sensory system globally, posing a significant public health issue that affects socio-political and economic aspects. However, whether and how NSD2, a histone methyltransferase, plays a role in HC maturation and hearing function are unclear. By HC-specific gene deletion of *Nsd2*, the present study demonstrates for the first time that NSD2 deficiency disrupts the epigenetic landscape of cochlear HCs to enhance the expression of ECM, and in turn impacts their maturation and hearing functions. This cell-autonomous pathological mechanism closely aligns with the clinical presentation of Wolf-Hirschhorn syndrome (WHS), thereby providing a novel molecular explanation for the sensorineural deafness observed in these patients. This discovery not only offers a novel perspective on the atypical role of NSD2, a new regulation factor of ECM, in the auditory system, but also paves a potential new path for uncovering the underlying mechanisms of hearing loss and cochlear dysfunction.

In view of that ECM is a vital component of the extracellular environment, providing structural support to cochlear HCs while also participating in cochlear development and sound conduction, the present work reveals an important cellular molecular mechanism as to how excessive ECM, especially collagen, mediates the hearing impairment caused by NSD2 deficiency. Firstly, we found that NSD2-deficient mice display impaired hearing function. (Fig. 2B). Secondly, histological examination of *Nsd2* mutant mice showed enlarged spaces between two adjacent HC bundles and increased width of the three rows of hair cells (Fig. 3A). Thirdly, single-cell sequencing revealed an increased expression of ECM associated genes, including *Egfl6, Mmp15, Sox9, Npnt, Col4a4*, *Ccdc80*. Fourthly, the upregulation of a key ECM protein collagen due to NSD2 deficiency was confirmed in HC-like HEI-OC1 cells. Interestingly, the excessive production of collagen appears to affect the area of HC-like HEI-OC1 cells on a substrate with a normal stiffness. Therefore, the overproduction of ECM induced by NSD2 deficiency may cause enhanced extracellular spaces and interfere their signal transmission, which leads to the hearing-impairment.

In support of our current study indicating that absence of NSD2 in cochlear HCs leads to a marked disruption of H3K36me2, previous studies indicate that NSD2 plays a pivotal role in regulating gene expression and maintaining cellular homeostasis and that the H3K36me2 modification exhibits intricate correlations with various other histone modifications such as H3K27me3 and H3K27ac, resulting in alterations to chromatin structure^43–46^. More importantly, current work shows that this disruption globally alters the chromatin accessibility to impact the transcriptional activities of genes related to hearing and ECM. In addition, the present RNA-seq and ATAC-seq experiments confirmed these abnormal transcription patterns of multiple genes related to ECM and HC maturation in NSD2-deficient HEI-OC1 cells, suggesting a crucial role for NSD2 in regulating the expression of the genes related to HC maturation by influencing epigenetic markers.

There is a limitation in our research in which *Nsd2* is deleted by using *Aoth1-Cre; Nsd2^f/f^* mice, in which *Nsd2* is deleted only after HC differentiation is initiated (in *Atoh1 (+)* cells). In the future, using *Sox2-Cre* or *p27-Cre* mice in which *Cre* is driven under the promoters of HC progenitor/supporting cell-specific genes will be better choices to achieve the goal to determine whether *Nsd2* affects sensory progenitors. And considering that the ATAC and CUT&Tag were performed in cell lines, the identical alterations in H3K36me2 and chromatin accessibility remain to be further characterized in vivo. Nevertheless, our present work sheds light on the current understanding of epigenetic impact on hearing functions, particularly with regard to the regulation of H3K36me2. Moreover, our study highlights the need for a more holistic approach to investigate histone methylation in hearing research, rather than focusing solely on individual enzymes or modification sites.

In conclusion, this study offers novel insights into the role of NSD2 in cochlear HCs and unravels the pathogenesis of hearing impairment caused by NSD2 deficiency. Our findings provide a promising new direction for treating Nsd2-deficiency-induced sensorineural hearing loss. Considering the recent clinical breakthroughs in AAV1-mediated gene therapies for deafness^47^, AAV-mediated knockdown of targeted ECM genes may offer a potential therapeutic strategy to reverse the auditory phenotype.

## Materials and Methods

### Mice

The *Atoh1^Cre^* mice (B6.Cg-Tg(*Atoh1-cre*)1Bfri/J, Strain #: 011104) were obtained from The Jackson Laboratory, and NSD2-floxed mice (*Nsd2^f/f^*) were kindly gifted by Prof. Jun Qin from Chinese Academy of Sciences. *Nsd2^f/f^*mice were mated with *Atoh1^Cre^* mice to generate *Atoh1^Cre^*; *Nsd2^f/+^* mice. *Atoh1^Cre^*; *Nsd2^f/+^*mice were mated with *Nsd2^f/f^* mice to generate HC-specific Nsd2 conditional knockout (*Atoh1-Cre; Nsd2^f/f^, hereafter designated as Nsd2^Atoh1-KO^)* mice. Littermate *Nsd2^f/f^*mice served as the control group. Both male and female mice were utilized across all experimental cohorts, with sample sizes comprising a minimum of n=5 biological replicates per group as specified. Preliminary statistical evaluations confirmed that sex had no significant main effect or interaction on any measured physiological parameters (p>0.05); therefore, data from male and female mice were pooled for final analysis. All mice were maintained under specific pathogen-free conditions with standardized humane endpoints. All experimental procedures were approved by the Animal Ethics Committee of School of Biomedical Engineering & Med-X Research Institute, Shanghai Jiao Tong University.

### AAV preparation and injection

To achieve targeted knockdown of Ogn and Col6a3 genes or ectopic delivery of Nsd2 in the sensory epithelium, we employed an adeno-associated virus serotype 2 (AAV2) vector platform, leveraging its natural cell-specific tropism for mammalian HC. AAVs were generated in HEK-293T cells co-transfected with pAAV[shRNA]-mCherry-U6, AAV2 and a helper plasmid. We used pAAV-CMV-mCherry-Nsd2(mouse), pAAV-CMV-mCherry-U6-sh-Ogn(mouse), pAAV-CMV-mCherry-U6-shCol6a3(mouse), pAAV-CMV-MCS-mCherry and pAAV-CMV-mCherry-U6-shNC. Collect the culture medium 48 hours after transfection. Lyse the cells by repeated freeze - thaw cycles, and then centrifuge at 4°C to collect the supernatant. AAVs were isolated via iodixanol gradient ultracentrifugation (140,000 g, 6 hours, 4°C). Finally, determined the titer of AAV by SYBR analysis (about 1*10^^11^ pfu/ml). Both male and female C57BL/6J mice at 2 weeks of age were anesthetized with tribromoethanol (500 mg/kg). To provide pre-operative analgesia, mice were subcutaneously administered with 2% meloxicam approximately 30 minutes prior to surgical incision. Shaved the hair behind the ear and made an incision of about 1 cm. Used dissecting forceps to perform blunt dissection on the subcutaneous tissue. Located the cochlear sac after tearing a part of the sternocleidomastoid muscle, and perforated it to expose the round window membrane. Injected about 1.5 μL of AAV into the inner ear through the round window. Sealed the round window membrane with tissue adhesive (3M, 1469SB) and sutured the wound. Mice recovered at 37°C.

### Cell culture

The House Ear Institute-organ of Corti 1 (HEI-OC1) auditory cell line (ATCC, RRID: CVCL_D899) was kindly gifted by Prof. Huawei Li from Fudan University. This cell line serves as a robust in vitro proxy representing an undifferentiated, relatively immature state of HC progenitors. Cells were cultured under 33 °C and 10% CO2 conditions in HyClone™ Dulbecco’s Modified Eagle Medium (DMEM)/ F12 1:1: Liquid (Cytiva, SH30023.01) containing 10% fetal bovine serum, without antibiotics. This cell line was free of mycoplasmal and viral contamination. The line was routinely screened and verified free of mycoplasmal and viral contamination. Stable partial knockout cells (*Nsd2 sg*) and corresponding control vector cells (*Cv2*) were generated via CRISPR/Cas9 editing. Control *Cv2* lines were verified to be comparable to unedited parental HEI-OC1 cells.

To evaluate biophysical cell responses, non-coated glass and standardized Matrigel substrates were engineered. The localized mechanical stiffness of the sensory epithelium was quantified using Atomic Force Microscopy (AFM) indentation on native neonatal mouse cochleae, establishing a range approximately from 1.05 to 1.25 kPa. In vitro hydrogel parameters were calibrated to a biophysical parity of approximately 1.25 kPa to accurately mimic the in vivo pericellular compliance.

For cell spreading analyses, “cell area” was defined as the total two-dimensional projected surface area occupied by an individual adhered cell body. This parameter was quantified across n=5 independent biological replicates tracking over 50 cells per experimental condition via automated border-detection algorithms using ImageJ software. To evaluate cell motility, Transwell migration assays were conducted. Transwell inserts containing porous membranes were seeded with 100,000 cells under normal stiffness conditions, and the absolute numbers of migrated cells were quantified 24 hours post-seeding across independent replicates to evaluate migratory capacity.

### RNA isolation and quantitative RT-qPCR

Total RNA was extracted from cultured either stable cell lines or freshly micro-dissected whole organ of Corti bulk tissues with an RNA extraction kit (BioTeke) following the manufacturer’s protocol. Reverse transcription was performed using a commercial RT kit (Takara) to generate cDNA. Quantitative real-time PCR (RT-qPCR) was subsequently conducted with TB Green real-time PCR analysis.

*Gapdh* served as the endogenous baseline control for data normalization. Relative gene expression folds were calculated using the 2^−ΔΔCt^ method. Data are presented as mean ± S.D. derived from a minimum of n=5 biological replicates per group. Student’s t-test was used to calculate the P value.

### Histology and IF staining

Mice at specific postnatal developmental stages (including P0, P2, P4, P7, P14, P21, P28, and 8 weeks) were euthanized via CO2 asphyxiation. To maintain spatial representation along the tonotopic axis, the cochleae were processed intact. For cross-sectional histology, intact temporal bones were fixed, decalcified in 10% EDTA, paraffin-embedded, and sectioned at a thickness of 5 um for classical Hematoxylin and Eosin (H&E) or Masson’s Trichrome staining to evaluate cross-sectional morphology, cellular spacing, and extracellular collagen deposition.

For whole-mount immunofluorescence (IF) surface preparations, the bony capsule and lateral wall were meticulously removed to isolate the intact sensory epithelium. Tissues were fixed in 4% paraformaldehyde (15 min, RT), permeabilized in 0.1% Triton X-100 in PBS (10 min, RT), and blocked in 10% FBS/PBS (20 min, RT). Primary antibodies including NSD2 (Abcam, Cat# ab75359), H3K36me2 (Abcam, Cat# ab9049), Collagen I (Abcam, Cat# ab21286), Collagen III (Abcam, Cat# EPR17673), Collagen IV (Abcam, Cat# ab6586), Collagen V (Abcam, Cat# ab7046), and the cell-type-specific hair cell markers Rabbit Anti-Myosin7a (Proteus Biosciences, Cat# 25-6790) and Mouse Anti-Myo7a (DSHB, Cat# MYO7A 138-1) were incubated overnight at 4°C. Following thrice washing with PBS, samples were incubated with Alexa Fluor 488-conjugated secondary antibodies (Jackson ImmunoResearch, Cat# 111-585-003), Alexa Fluor 594 Goat Anti-Mouse IgG (H+L) (Jackson ImmunoResearch, Cat# 115-585-003, RRID: AB_2338871), and Phalloidin-Alexa Fluor 647 (Thermo Fisher Scientific, Cat# A22287) for 2 hours at room temperature. Finally, the specimens were mounted using an anti-fade mounting medium containing DAPI (Vector Laboratories, Cat# H-1200-10) for nuclear counterstaining and structural preservation.

Laser confocal microscopy (Leica) was implemented for high-resolution imaging. To eliminate vertical topography variations, optical scanning parameters including laser intensity, photomultiplier gain, and pinhole diameter were held fully constant across all parallel groups. Semi-quantitative fluorescence analyses were strictly conducted on single horizontal optical planes centered at the reticular lamina level. Regions of Interest (ROIs) were defined by manually outlining individual sensory cell somas based on the boundary delineated by the Myo7a signal. Mean fluorescence intensity values were quantified using ImageJ software by subtracting background noise across a minimum of 20 individual HC per turn across n = 5 independent biological samples. Anatomical metrics, including outer hair cell (OHC) spacing and the total width covering the three rows of OHC, were measured across the distinct apical, middle, and basal turns. Data were presented as grouped box plots and analyzed via ANOVA with Tukey’s post hoc test.

### ABR Measurement

Auditory thresholds and electrophysiological waveforms were recorded within a specialized sound-attenuating, radiofrequency-shielded booth (Shanghai Shino Acoustic Equipment Co., Ltd). Mice were anesthetized via continuous isoflurane inhalation and stabilized on a feedback-controlled heating pad to maintain core normothermia. Subdermal needle electrodes were positioned at the vertex (active), right mastoid (reference), and left shoulder (ground). Detailed assay protocols are described in a previous report^48^. Acoustic stimuli comprised both brief click transients and specific tone-burst frequencies (4, 5.6, 8, 11.3, 16, 22.6, and 32 kHz) generated by a digital signal processor.

To eliminate threshold-shift artifacts on propagation metrics, ABR waveform reproducibility, wave amplitudes (I to VI), and absolute peak latencies were systematically extracted and quantified from a stabilized suprathreshold stimulus fixed precisely at 20 dB SPL above each individual animal’s determined hearing threshold. ABR waveforms present individual animal recordings alongside the calculated group grand average represented by a stabilized solid trace. Statistical differences between groups were assessed via Student’s t-test (p<0.05).

### Single-cell RNA Sequencing

The NovelCyto Single-Cell Analysis System (NovelBio Co., Ltd., China) was implemented for high-throughput transcriptome capture. Single-cell suspensions were stochastically partitioned into >100,000 microwells via limited dilution. Oligonucleotide-coupled magnetic beads, pre-loaded with unique barcodes, were introduced until saturation, achieving that each bead paired with a cell in a microwell. Following lysis, polyadenylated mRNA molecules were captured through hybridization with bead-barcoded capture oligos. Beads were subsequently pooled for reverse transcription and ExoI digestion. During cDNA synthesis, each transcript was labeled at the 5′-terminus with a unique molecular identifier (UMI) and cell barcode, enabling precise quantification of mRNA copies and unambiguous cell-of-origin tracking.

Whole-transcriptome libraries were constructed using the NovelCyto single-cell WTA workflow, comprising random hexamer priming/extension, PCR amplification, and library indexing PCR. Library quality was assessed via a High Sensitivity DNA chip (Agilent) on a Bioanalyzer 4200 and a Qubit High Sensitivity DNA assay (Thermo Fisher Scientific). Final libraries were sequenced on an Illumina NovaSeq 6000 platform (Illumina, San Diego, CA) using a paired-end 150 bp (PE150) configuration.

### RNA sequencing and analyses

Total RNA was isolated using TRIzol (Invitrogen), with RNA integrity number (RIN >7.0) via Agilent 2200 Bioanalyzer. Stranded mRNA libraries were prepared using the Illumina TruSeq Stranded mRNA Library Prep Kit. Sequencing reads were quality-filtered (adapters/low-quality bases removed) and aligned to the mouse genome (mm10) using HISAT2. Gene counts were quantified via HTSeq, with expression levels normalized as RPKM. The R package of edgeR v3.24.2 was used to measure differential gene expression. Differential expression analysis (|log2FC|>1, adj.P<0.05) was conducted using DESeq2. Gene annotations were retrieved from the Ensembl genome browser 96 database (http://www.ensembl.org/index.html), and functional enrichment analysis of differentially expressed genes (DEGs) for Gene Ontology (GO) terms and KEGG pathways was performed using the ClusterProfiler R package, integrating annotations from NCBI, UniProt, and the GO Consortium. Statistical significance of enriched terms/pathways was determined by Fisher’s exact test (P<0.05).

### CUT&Tag assay and analyses

CUT&Tag was performed using the Hyperactive® Universal CUT&Tag Assay Kit for Illumina (Vazyme, TD903) with nuclei isolated from 100,000 fresh cells. Nuclei were bound to ConA beads, incubated with primary antibody (H3K36me2 [Abcam ab9049] H3K36me2 [Abcam ab9049]) or IgG control, followed by secondary antibody and pA-Tn5 adapter complex (1 h, RT). DNA was fragmented with 5×TTBL, purified via proteinase K digestion and magnetic beads, and libraries amplified with 14 PCR cycles. Sequencing was conducted on an Illumina NovaSeq 6000 with paired-end 2x150 as the sequencing mode. Raw reads were processed using Trimmomatic v0.38 (SLIDINGWINDOW:4:15, LEADING:10, TRAILING:10, MINLEN:36) to remove adapters, short/low-quality reads.

FastQC confirmed read quality. Clean reads were aligned to GRCm38 with Bowtie2 v2.3.4.1 (-I 10 -X 700 --no-discordant --no-mixed --local --very-sensitive-local). Duplicates were removed via Picard MarkDuplicates. Peaks were called using MACS2 v2.1.2 (-f BAMPE -g mm -q 0.05) and annotated with ChIPseeker R package.

This method leverages the high activity of Tn5 transposase fused to Protein A/G (pA/G-Tn5), which binds to antibody-labeled chromatin regions under native conditions. Upon binding, the Tn5 enzyme simultaneously fragments the DNA (Cut) and ligates sequencing adapters to both ends of the cleaved fragments. The resulting DNA fragments are then PCR-amplified to generate sequencing-ready libraries (Tag).

Data are presented as mean ± SEM (n=3 – 15 mice or ≥ 3 replicates). Statistical significance was assessed using Student’s t-test or two-way ANOVA (GraphPad 8.0), with significance thresholds set at *p<0.05, **p<0.01, ***p<0.001, ****p<0.0001.

### ATAC- sequencing and analyses

ATAC-seq was performed on 50,000 cells lysed in cold buffer (10 mM Tris-HCl pH 7.4, 10 mM NaCl, 3 mM MgCl₂, 0.1% IGEPAL CA-630). Nuclei were pelleted (500 ×g, 10 min, 4°C) and tagged using Tn5 transposase (Illumina) in TD buffer (37°C, 30 min). Libraries were purified (Qiagen MinElute), amplified with NEBNext PCR Master Mix and Nextera primers (72°C/5 min; 98°C/30 s; 12 cycles of 98°C/10 s, 63°C/30 s, 72°C/1 min), and sequenced (Illumina).

Raw reads were trimmed (Trimmomatic v0.38: SLIDINGWINDOW:4:15, LEADING/TRAILING:10, MINLEN:36) and quality-checked (FastQC). Clean reads were aligned to mm10 (Bowtie2 v2.3.4.1: -N 1 -L 25 -X 2000 --no-mixed --no-discordant), filtered (SAMtools MAPQ≥30), and deduplicated (Picard). Peaks were called using MACS2 v2.1.2 (-g mm -q 0.05 --nomodel) and filtered via IDR (v2.0.2, IDR≤0.05). ChIPseeker R package annotated peaks to genomic features, while MEME-suite identified motifs. Differential peaks (DiffBind, q<0.05) and functional enrichment (ClusterProfiler v3.4.4, GO/KEGG) were analyzed.

### Statistical Analysis

All data are presented as mean ± SEM or mean ± S.D. as specified, derived from a minimum of *n* = 5 or *n* = 6 biological mouse samples or at least three independent biological cell line replicates. Statistical evaluations were performed using GraphPad Prism 8.0 software. Group comparisons were conducted using an unpaired two-tailed Student’s *t* -test or a two-way ANOVA followed by Tukey’s post hoc test. Statistical significance thresholds were defined directly within the relevant graphs as ∗ *P* < 0.05, ∗∗ *P* < 0.01, ∗∗∗ *P* < 0.001, and ∗∗∗∗ *P* < 0.0001. All raw sequencing data and processed metadata hubs have been deposited into the NCBI Gene Expression Omnibus (GEO) public repository under secure accession tokens. The clean reads of RNA-seq were then aligned to mouse genome (version: GRCm38 NCBI) using the HISAT2 v2.2.1.Then CUT&Tag reads were aligned to remove adapters, short/low-quality reads using Trimmomatic (v0.38) software, remaining reads were used to align to reference genome and transcriptome (GRCm38) using MACS2 (v2.1.2) and BOWTIE2 (v2.3.4.1) software separately. Then ATAC-seq reads were aligned to remove adapters, short/low-quality reads using Trimmomatic (v0.38) software, remaining reads were used to align to reference genome and transcriptome (mm10) using MACS2 (v2.1.2) and IDR (v2.0.2) software separately.DIA-NN (version 1.8.1) was used to collect the MS data.Differential peaks and functional enrichment (ClusterProfiler v3.4.4) were analyzed.

## Acknowledgments

This work was supported by funds from National Natural Science Foundation of China (U23A20441, W2431055, 82573626 to W.-Q.G., 32570684, 82372604 to L.L.), National Key R&D Program of China (2022YFA1302704 to L.L. and W.-Q.G.), the Peak Disciplines (Type IV) of Institutions of Higher Learning in Shanghai, 111 project (B21024) and KC Wong foundation to W.-Q.G and Interdisciplinary Program of Shanghai Jiao Tong University (YG2024ZD11). We thank Genefund Biotech (Shanghai, China) and NovelBio (Shanghai, China) for assistance in the data analysis. We thank Qiao Yu and Prof. Hao He (Shanghai Jiao Tong University) for providing access to confocal microscopy facilities.

## Author contributions

L.L. and W.-Q.G. conceived the experimental concept, designed the experiments and interpreted the data; Z.W. performed most of the experiments and wrote the manuscript; X.M. and Y.X. helped with the clinical data analysis; D.W., W.F., H.R., W.Z., N.L., R.A., J.L., G.Y. assisted in some experiments; W.-Q.G. assisted in some discussion; Z.W. wrote the manuscript; L.L., provided the overall guidance. All authors read and approved the final manuscript.

## Competing interests

The authors have declared that no conflict of interest exists.

## Data Availability Statement

The authors declare that all data supporting the findings in this study are available within the paper, Supplementary information, and Source data. All the raw data have been deposited in the Gene Expression Omnibus (GEO) under accession number GEO: GSE282857(Single-cell RNA-Seq), GSE281335 (RNA-Seq), GSE281336 (CUT-Tag), GSE281334 (ATAC-Seq). All data are available from the authors upon reasonable request.

## Notes

### Competing Interest Statement

The authors have declared no competing interest.

## References

1. Géléoc, G. S. G. & Holt, J. R. Sound Strategies for Hearing Restoration. Science 344, 1241062 (2014).

2. Laszig, R. & Aschendorff, A. Cochlear implants and electrical brainstem stimulation in sensorineural hearing loss. Curr Opin Neurol 12, 41–44 (1999).

3. Weisleder, P. & Rubel, E. W. Hair cell regeneration after streptomycin toxicity in the avian vestibular epithelium. J Comp Neurol 331, 97–110 (1993).

4. Jennifer S Stone and Douglas Allen Cotanche. [PDF] Hair cell regeneration in the avian auditory epithelium. | Semantic Scholar. https://www.semanticscholar.org/paper/Hair-cell-regeneration-in-the-avian-auditory-Stone-Cotanche/f555cd06f5db89747512a3a6fce6cca9e127d71a.

5. Corwin, J. T. & Cotanche, D. A. Regeneration of sensory hair cells after acoustic trauma. Science 240, 1772–1774 (1988).

6. Zheng, J. L. & Gao, W. Q. Overexpression of Math1 induces robust production of extra hair cells in postnatal rat inner ears. Nat Neurosci 3, 580–586 (2000).

7. Zheng, J. L., Shou, J., Guillemot, F., Kageyama, R. & Gao, W. Q. Hes1 is a negative regulator of inner ear hair cell differentiation. Development 127, 4551–4560 (2000).

8. Chen, Y. et al. Hedgehog signaling promotes the proliferation and subsequent hair cell formation of progenitor cells in the neonatal mouse cochlea. Frontiers in Molecular Neuroscience 10, 426 (2017).

9. Chen, Q. et al. Inactivation of STAT3 Signaling Impairs Hair Cell Differentiation in the Developing Mouse Cochlea. Stem Cell Reports 9, 231–246 (2017).

10. Ni, W. et al. Wnt activation followed by Notch inhibition promotes mitotic hair cell regeneration in the postnatal mouse cochlea. Oncotarget 7, 66754–66768 (2016).

11. Waqas, M., Zhang, S., He, Z., Tang, M. & Chai, R. Role of Wnt and Notch signaling in regulating hair cell regeneration in the cochlea. Front Med 10, 237–249 (2016).

12. Pressé, M. T., Malgrange, B. & Delacroix, L. The cochlear matrisome: Importance in hearing and deafness. Matrix Biol 125, 40–58 (2024).

13. Tsuprun, V. & Santi, P. Proteoglycan arrays in the cochlear basement membrane. Hear Res 157, 65–76 (2001).

14. Thalmann, I. Collagen of accessory structures of organ of corti. Connective Tissue Research 10.3109/03008209309016826 (1993) doi:10.3109/03008209309016826.

15. Thalmann, I., Thallinger, G., Comegys, T. H. & Thalmann, R. Collagen – The Predominant Protein of the Tectorial Membrane. ORL 48, 107–115 (2010).

16. Xia, M. et al. Varying mechanical forces drive sensory epithelium formation. Sci Adv 9, eadf2664 (2023).

17. Amma, L. L. et al. An emilin family extracellular matrix protein identified in the cochlear basilar membrane. Mol Cell Neurosci 23, 460–472 (2003).

18. Chen, J. et al. Single-Cell RNA Sequencing Analysis Reveals Greater Epithelial Ridge Cells Degeneration During Postnatal Development of Cochlea in Rats. Front Cell Dev Biol 9, 719491 (2021).

19. Mittal, R. et al. Recent advancements in understanding the role of epigenetics in the auditory system. Gene 761, 144996 (2020).

20. Mittal, R. et al. Recent advancements in understanding the role of epigenetics in the auditory system. Gene 761, 144996 (2020).

21. Shin, J.-O. et al. CTCF Regulates Otic Neurogenesis via Histone Modification in the Neurog1 Locus. Mol Cells 41, 695–702 (2018).

22. Stojanova, Z. P., Kwan, T. & Segil, N. Epigenetic regulation of Atoh1 guides hair cell development in the mammalian cochlea. Development 142, 3529–3536 (2015).

23. Patel, D., Shimomura, A., Majumdar, S., Holley, M. C. & Hashino, E. The histone demethylase LSD1 regulates inner ear progenitor differentiation through interactions with Pax2 and the NuRD repressor complex. PLoS One 13, e0191689 (2018).

24. Ahmed, M. & Streit, A. Lsd1 interacts with cMyb to demethylate repressive histone marks and maintain inner ear progenitor identity. Development 145, dev160325 (2018).

25. He, L., Cao, Y. & Sun, L. NSD family proteins: Rising stars as therapeutic targets. Cell Insight 3, 100151 (2024).

26. Bennett, R. L., Swaroop, A., Troche, C. & Licht, J. D. The Role of Nuclear Receptor–Binding SET Domain Family Histone Lysine Methyltransferases in Cancer. Cold Spring Harb Perspect Med 7, a026708 (2017).

27. Yee, S. P. & Rigby, P. W. The regulation of myogenin gene expression during the embryonic development of the mouse. Genes Dev. 7, 1277–1289 (1993).

28. Bennett, R. L., Swaroop, A., Troche, C. & Licht, J. D. The Role of Nuclear Receptor–Binding SET Domain Family Histone Lysine Methyltransferases in Cancer. Cold Spring Harb Perspect Med 7, a026708 (2017).

29. Zanoni, P. et al. Loss-of-function and missense variants in NSD2 cause decreased methylation activity and are associated with a distinct developmental phenotype. Genet Med 23, 1474–1483 (2021).

30. Stec, I. et al. WHSC1, a 90 kb SET Domain-Containing Gene, Expressed in Early Development and Homologous to a Drosophila Dysmorphy Gene Maps in the Wolf-Hirschhorn Syndrome Critical Region and is Fused to IgH in t(1;14) Multiple Myeloma. Human Molecular Genetics 7, 1071–1082 (1998).

31. Nimura, K. et al. A histone H3 lysine 36 trimethyltransferase links Nkx2-5 to Wolf-Hirschhorn syndrome. Nature 460, 287–291 (2009).

32. Ahmed, M., Ura, K. & Streit, A. Auditory hair cell defects as potential cause for sensorineural deafness in Wolf-Hirschhorn syndrome. Disease Models & Mechanisms 8, 1027–1035 (2015).

33. Li, S. et al. Fate-mapping analysis of cochlear cells expressing Atoh1 mRNA via a new Atoh13*HA-P2A-Cre knockin mouse strain. Dev Dyn 251, 1156–1174 (2022).

34. Li, N. et al. AKT-mediated stabilization of histone methyltransferase WHSC1 promotes prostate cancer metastasis. J Clin Invest 127, 1284–1302 (2017).

35. Matei, V. et al. Smaller inner ear sensory epithelia in Neurog1 null mice are related to earlier hair cell cycle exit. Developmental Dynamics 234, 633–650 (2005).

36. Zaaroor, M. & Starr, A. Auditory brain-stem evoked potentials in cat after kainic acid induced neuronal loss. II. Cochlear nucleus. Electroencephalography and Clinical Neurophysiology/Evoked Potentials Section 80, 436–445 (1991).

37. Jewett, D. L., Romano, M. N. & Williston, J. S. Human Auditory Evoked Potentials: Possible Brain Stem Components Detected on the Scalp. Science 167, 1517–1518 (1970).

38. Kalinec, G. M., Webster, P., Lim, D. J. & Kalinec, F. A cochlear cell line as an in vitro system for drug ototoxicity screening. Audiology and Neuro-Otology 8, 177–189 (2003).

39. Devarajan, P. et al. Cisplatin-induced apoptosis in auditory cells: Role of death receptor and mitochondrial pathways. Hearing Research 174, 45–54 (2002).

40. Li, Z. et al. H3K36me2 methyltransferase NSD2 orchestrates epigenetic reprogramming during spermatogenesis. Nucleic Acids Res 50, 6786–6800 (2022).

41. Kinoshita, S. et al. Loss of NSD2 causes dysregulation of synaptic genes and altered H3K36 dimethylation in mice. Front. Genet. 15, (2024).

42. Zhuang, L. et al. Depletion of Nsd2-mediated histone H3K36 methylation impairs adipose tissue development and function. Nat Commun 9, 1796 (2018).

43. Fang, Y. et al. The H3K36me2 methyltransferase NSD1 modulates H3K27ac at active enhancers to safeguard gene expression. Nucleic Acids Res 49, 6281–6295 (2021).

44. Chen, R. et al. Di- and tri-methylation of histone H3K36 play distinct roles in DNA double-strand break repair. Sci China Life Sci 67, 1089–1105 (2024).

45. Li, Y. et al. Histone methylation antagonism drives tumor immune evasion in squamous cell carcinomas. Mol Cell 82, 3901–3918.e7 (2022).

46. Sun, Z. et al. Chromatin regulation of transcriptional enhancers and cell fate by the Sotos syndrome gene NSD1. Mol Cell 83, 2398–2416.e12 (2023).

47. Lv, J. et al. AAV1-hOTOF gene therapy for autosomal recessive deafness 9: a single-arm trial. The Lancet 403, 2317–2325 (2024).

48. Li, C. et al. Characterizing a novel vGlut3-P2A-iCreER knockin mouse strain in cochlea. Hear Res 364, 12–24 (2018).

