## supplemental figure for "NSD2 deficiency disrupts the epigenetic landscape of cochlear hair cells to enhance ECM expression and to perturb cochlear hair cell function"

**Fig. S1.**

**(A)** Relative expression levels of Nsd2 in isolated cochlear hair cells at different developmental stages (E16, P0, P4, and P7). Data were analyzed using the public dataset GSE60019. Data are presented as relative expression calculated by the $2^{-\Delta\Delta CT}$ method.

**(B)** Representative agarose gel images showing the identification of Atoh1-cre (left) and floxed Nsd2 (right) alleles, with analytical samples (AN), positive (+), and negative (-) controls cropped from at least three independent experimental replicates to ensure technical reproducibility.

**(C)** RT-qPCR analysis (as shown in image_09a6da.png) shows a significant reduction of Nsd2 mRNA expression levels in the Organ of Corti of *Nsd2*$^{f/f}$ and *Nsd2*$^{Atoh1\text{-}KO}$ mice (n = 5 mice per group), with relative expression calculated using the $2^{-\Delta\Delta CT}$ method and data presented as mean ± SEM; *P < 0.05 by Student's t-test.

**(D)** Representative whole-mount immunofluorescence images (as shown in image_0a9797.jpg) and statistical analysis of the total width covering the three rows of outer hair cells (OHCs) in the basal (top) and apical (bottom) turns of *Nsd2*$^{f/f}$ and *Nsd2*$^{Atoh1\text{-}KO}$ mice. Cochleae were immunostained with antibodies against Myo7a (green) to mark hair cells, phalloidin (red) to visualize F-actin, and DAPI (blue) for nuclear counterstaining. To eliminate focal depth artifacts, all quantified images were captured within perfectly matched focal planes by averaging 3–4 consecutive z-stack sections. Quantitative analysis confirmed a significant expansion of the OHC row width in Nsd2f/f and Nsd2Atoh1-KO mice. Data are presented as mean ± SEM (n = 5 mice per group); *P < 0.05 by Student's t-test.Staining of Myo7a in cochleae base and apical turns of adult *Nsd2*$^{f/f}$ and *Nsd2*$^{Atoh1\text{-}KO}$ mice and quantification of Myo7a fluorescence intensity (n = 5 per group).

**Fig. S2.**

**(A)**. Statistical analysis quantifying general hair cell (HC) density per 100 $\mu$m (left) and the individual spacing between outer hair cells ($\mu$m, right) in the cochleae of $Nsd2^{f/f}$ and $Nsd2^{Atoh1\text{-}KO}$ mice. While overall HC density remains unaltered (ns, non-significant), a significant increase in outer hair cell spacing is observed upon *Nsd2* deficiency. Data are presented as mean $\pm$ SEM ($n = 5$ mice per group); $** P < 0.01$ by Student's $t$-test

**(B)** Statistical analysis demonstrating spatial and structural alterations in the cochleae of $Nsd2^{\text{f/f}}$ and $Nsd2^{\text{Atoh1-KO}}$ mice. Quantitative parameters include the total width covering 3 rows of OHCs ($\mu$m, left), OHC stereocilia inter-bundle distance ($\mu$m, middle), and total hair cell (HC) density per 68 $\mu$m (right). *Nsd2* deficiency significantly increases both the overall row width and the inter-bundle distance between neighboring stereocilia, whereas total hair cell numbers remain unchanged (ns, non-significant). Data are presented as mean $\pm$ SEM ($n = 5$ mice per group); $* P < 0.05$, $** P < 0.01$ by Student's $t$-test.

**(C)** RT-qPCR analysis (as shown in image_0c68f8.png) showing the relative mRNA expression levels of stereocilia- and hearing-related markers—including *Myo3a*, *Tmc1*, *Espnl*, *Tomt*, and *Gjb2*—in the Organ of Corti of $Nsd2^{\text{f/f}}$ and $Nsd2^{\text{Atoh1-KO}}$ mice. Conditional deletion of *Nsd2* leads to a significant downregulation of all tested functional target genes. Relative expression was calculated using the $2^{-\Delta\Delta C_T}$ method, and data are presented as mean $\pm$ SEM ($n = 5$ mice per group); $* P < 0.05$, $** P < 0.01$, $*** P < 0.001$ by Student's $t$-test.

**(D)** Heatmaps displaying representative differentially expressed genes (DEGs) clustered by key Gene Ontology (GO) terms—including *Autophagic cell death and positive regulation of apoptotic process*, *Negative regulation of epithelial cell proliferation*, *Extracellular matrix organization*, *Cell differentiation*, *Regulation of cell cycle*, and *Nervous system development*—comparing control ($Nsd2^{\text{f/f}}$) and knockout ($Nsd2^{\text{f/f}}; Atoh1$-cre) groups. Expression levels are represented by z-scores (blue, downregulated; orange, upregulated).

**(E)** RT-qPCR validation of representative target genes associated with *Extracellular matrix organization* (*Ccdc80*, *Egfl6*, *Mmp15*, *Col4a4*, *Sox9*, and *Npnt*) in the Organ of Corti of $Nsd2^{\text{f/f}}$ and $Nsd2^{\text{Atoh1-KO}}$ mice. Relative expression was calculated using the $2^{-\Delta\Delta C_T}$ method. Data are presented as mean $\pm$ SEM ($n = 4$ mice per group); ns, non-significant; $* P < 0.05$, $*** P < 0.001$, $**** P < 0.0001$ by Student's $t$-test.

**Fig. S3.**

**(A-B)** Representative immunofluorescence panels and statistical quantification demonstrating a highly significant reduction in NSD2 protein levels (green) in *Nsd2* Sg cells compared to Cv2 controls ($n = 4$ independent cell culture samples per group). Nuclei were counterstained with DAPI (blue). Scale bar, 50 $\mu$m.

**(C-D)** Representative immunofluorescence panels and statistical quantification showing a concomitant decrease in histone H3K36me2 intensity (green) following Nsd2 depletion (n = 5 independent cell culture samples per group). Scale bar, 50 $\mu$m

**(E)** RT-qPCR analysis of functional hair cell-related and developmental marker genes (*Myo3a*, *Tmc1*, *Nsd2*, *Notch1*, *Espnl*, and *Tomt*) following *Nsd2* knockdown ($n = 5$ independent technical replicates per group)

**(F)** RT-qPCR profiling validating the marked upregulation of key extracellular matrix (ECM) components and organizational genes (*Bcam*, *Npnt*, *Tecta*, *Ccdc80*, and *Col4a4*) in *Nsd2* Sg HEI-OC1 cells ($n = 4$ independent technical replicates per group). Relative expression levels were normalized via the $2^{-\Delta\Delta C_T}$ method. All data are presented as mean $\pm$ SEM; $*P < 0.05$, $**P < 0.01$, $****P < 0.0001$ by Student's $t$-test.

**Fig. S4.**

**(A)** Representative whole-mount immunofluorescence images (as shown in image_0ce0f6.jpg) and intensity quantification demonstrating the deposition of structural collagens—Collagen 4, Collagen 5 (co-labeled with phalloidin, red), and Collagen 1 (co-labeled with Myo7a, red)—counterstained with DAPI (blue) within the hair cell region of control ($Nsd2^{\text{f/f}}$) and knockout ($Nsd2^{\text{Atoh1-KO}}$) mice. Conditional deletion of *Nsd2* leads to a pronounced accumulation of all tested extracellular matrix collagens surrounding the outer hair cells. To eliminate focal depth artifacts, fluorescence intensities were quantified from standardized regions across matched z-stack sections, with data presented as mean $\pm$ SEM ($n = 5$ mice per group); $*P < 0.05$, $***P < 0.001$ by Student's $t$-test.

**Fig. S5.**

**(A)** RT-qPCR validation demonstrating significant knockdown efficiency of *Ogn* and *Col6a3* mRNA levels following AAV-shRNA delivery ($n = 5$ mice per group).

**(B-C)** Morphometric analysis showing that AAV-mediated knockdown of either *Ogn* or *Col6a3* significantly rescues the abnormal expansion of the three-row OHC width ($\mu$m) and individual outer hair cell spacing ($\mu$m) observed in $Nsd2^{\text{Atoh1-KO}}$ mice back toward control levels ($n = 5$ mice per group).

**(C)** RT-qPCR validation confirming robust overexpression of *Nsd2* mRNA levels in the Organ of Corti following *Nsd2*-AAV virus delivery ($n = 5$ mice per group).

**(D)** Morphometric analysis demonstrating that exogenous rescue via *Nsd2*-AAV significantly reduces both the total width covering 3 OHCs ($\mu$m) and the expanded outer hair cell spacing ($\mu$m) in $Nsd2^{\text{Atoh1-KO}}$ mice ($n = 5$ mice per group).

**Fig. S6.**

**(A-C)** Morphological analysis and cell area quantification ($\mu$m$^2$) of Cv2 and *Nsd2* sg HEI-OC1 cells cultured on hydrogels of varying stiffness (0.42 kPa, 0.65 kPa, 0.97 kPa, and 1.25 kPa) at Day 1 **(A, B)** and Day 2 **(A, C)**, demonstrating that *Nsd2* deficiency significantly suppresses stiffness-dependent cell spreading ($n = 4$--$5$ independent samples per group). Scale bars, 100 $\mu$m.

**(D)** Representative Transwell migration assay images and statistical quantification of migrated cell numbers of $Nsd2^{\text{f/f}}$ and $Nsd2^{\text{Atoh1-KO}}$ cells across soft (0.42 kPa) and stiff (1.25 kPa) environments ($n = 5$ independent experiments), showing altered migratory capacity sensitive to substrate mechanics.

All data are presented as mean $\pm$ SEM; $*P < 0.05$, $**P < 0.01$, $***P < 0.001$, $****$ $P < 0.0001$ by Student's $t$-test or two-way ANOVA.

**Fig. S7.**

**(A)** Representative cell morphology images and statistical quantification of cell area ($\mu$m$^2$)

in control (Blank) and Nsd2-overexpressing (Nsd2 oe) cells cultured on soft (0.42 kPa) and stiff (0.97 kPa) hydrogels ($\mathrm{n} = 5$ independent samples per group), showing that exogenous Nsd2 expression significantly shifts cell spreading characteristics depending on matrix mechanics. Scale bar, 100 $\mu$m.

**(B)** RT-qPCR analysis tracking the transcriptional changes of major extracellular matrix components—*Col4a4*, *Ogn*, and *Col3a1*—across distinct matrix stiffness environments (0.42 kPa, 0.65 kPa, and 0.97 kPa) upon *Nsd2* knockdown (*Nsd2* sg vs. Cv2) or *Nsd2* overexpression (*Nsd2* oe vs. Blank) ($n = 5$ independent technical replicates per group). The data demonstrate that *Nsd2* deficiency consistently accentuates, while *Nsd2* overexpression represses, the expression of these mechanically responsive ECM genes. Relative expression levels were determined using the $2^{-\Delta\Delta C_T}$ method. Data are expressed as mean $\pm$ SEM; $* P < 0.05$, $** P < 0.01$, $*** P < 0.001$, $**** P < 0.0001$ by Student's $t$-test or two-way ANOVA.

**Fig. S8.**

**(A)** Representative cell morphology images on Day 1 (left) and Day 2 (right) of *Nsd2* sg, *Nsd2* sg *Ogn*-KO, *Nsd2* sg *Col6a3*-KO, and Cv2 control cells cultured on ultra-soft hydrogels (50 Pa and 30.8 Pa), showing structural adaptations. Scale bars, 100 $\mu$m.

**(B)** Statistical quantification of cell area ($\mu$m$^2$) on Day 2 across different substrate stiffnesses (1.25 kPa and 0.97 kPa). Knockout of downstream ECM genes (*Ogn* or *Col6a3*) significantly rescues the abnormal cell spreading phenotypes induced by *Nsd2* deficiency ($n = 5$ independent samples per group).

**(C)** RT-qPCR validation confirming the targeted knockout efficiency of Ogn and Col6a3, alongside accompanying changes in Ccdc80 and Col3a1 mRNA expression levels in control (Cv2) and Nsd2-depleted (Sg2) backgrounds ($\mathrm{n} = 5$ independent technical replicates per group).

**(D)** RT-qPCR analysis demonstrating that genetic ablation of *Ogn* or *Col6a3* in *Nsd2* sg cells significantly rescues the expression of key functional hair cell and developmental markers (*Myo3a*, *Tmc1*, *Nsd2*, *Notch1*, *Espnl*, and *Tomt*) on a 1.25 kPa stiff substrate at

Day 2 ($n = 5$ independent technical replicates per group).

Relative expression levels were determined using the $2^{-\Delta\Delta C_T}$ method. Data are expressed as mean $\pm$ SEM; $*P < 0.05$, $**P < 0.01$, $***P < 0.001$, $****P < 0.0001$ by Student's $t$-test or ANOVA.

**Fig. S9.**

**(A)** Hierarchical clustering heatmap of differentially expressed genes (DEGs) displaying distinct transcriptional profiles between the cv2 and *Nsd2* sg groups (color bar indicates z-score).

**(B)** Volcano plot showing significantly upregulated (red; 118 genes) and downregulated (blue; 325 genes) DEGs, with unchanged genes in gray (no-DEGs; 15,041 genes) based on $\log_2$(fold change) and $-\log_{10}(q$-value).

**(C)** Functional enrichment analysis of RNA-seq data outlining significantly altered pathways and structural terms, highlighting prominent changes in *extracellular matrix* and *collagen-containing extracellular matrix* (red box), alongside histone, chromatin, and hair cell-related terms. Numbers represent enrichment scores ($-\log_{10} P$-value or fold enrichment).

**(D)** Gene Set Enrichment Analysis (GSEA) plots demonstrating significant enrichment of genes associated with *Structural constituent of chromatin* (NES $= 1.942$, $P < 0.001$) and *Protein DNA complex assembly* (NES $= 1.651$, $P < 0.001$) in *Nsd2*-deficient cells

**(E)** Hierarchical clustering heatmaps of representative DEGs involved in specific enriched biological processes, including *Negative regulation of neuron death*, *Negative regulation of cell development*, *Negative regulation of gene expression, epigenetic*, and *Epidermal cell differentiation* (including *Myo7a*). Color scaling denotes relative gene expression (blue, downregulated; orange/red, upregulated).

**Fig. S10.**

**(A)** Transcription factor motif enrichment analysis within regions of differentially accessible chromatin. The bar plot indicates significantly enriched motifs in regions of open chromatin (red, upregulated accessibility) and closed chromatin (blue, downregulated accessibility) based on $-\log_{10}(P\text{-value})$.

**(B)** Representative ATAC-seq genomic tracks and corresponding RNA-seq expression levels at the *Ccdc80*, *Col3a1*, and *Igf1* genomic loci in control ($Nsd2^{\text{f/f}}$, blue) and knockout ($Nsd2^{\text{Atoh1-KO}}$, red) groups. Light pink shaded areas highlight key transcriptionally active regions or regulatory elements where alterations in chromatin accessibility correspond with changes in mRNA expression level.

Fig.S1

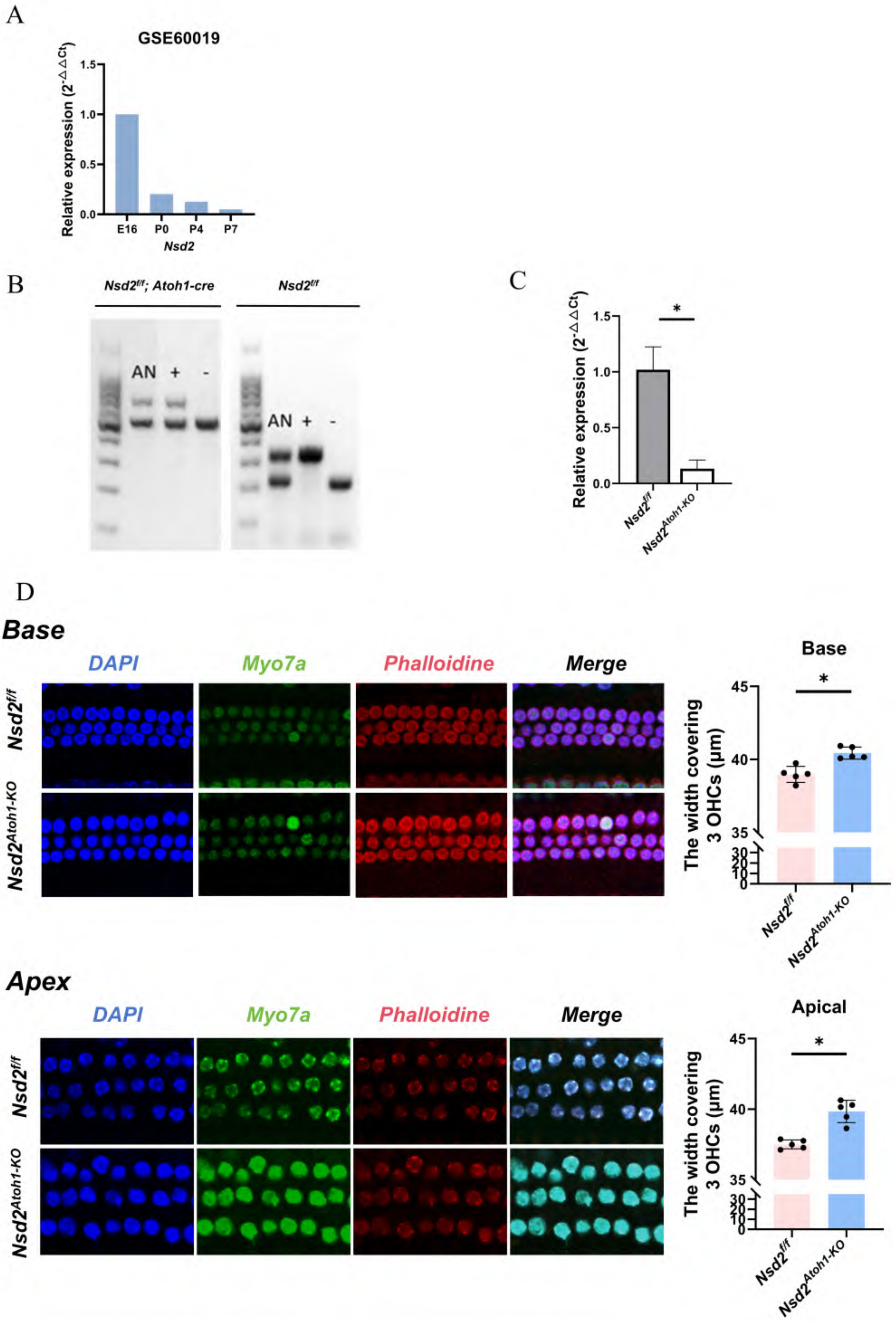
A
GSE60019
Relative expression (2-ΔΔCt)
1.5
1.0
0.5
0.0
E16
P0
P4
P7
Nsd2
B
Nsd2f/f; Atoh1-cre
Nsd2f/f
AN
+
-
C
*
Nsd2f/f
Nsd2Atoh1-KO
D
Base
DAPI
Myo7a
Phalloidine
Merge
The width covering 3 OHCs (μm)
45
40
35
Apex
Apical

Fig.S2

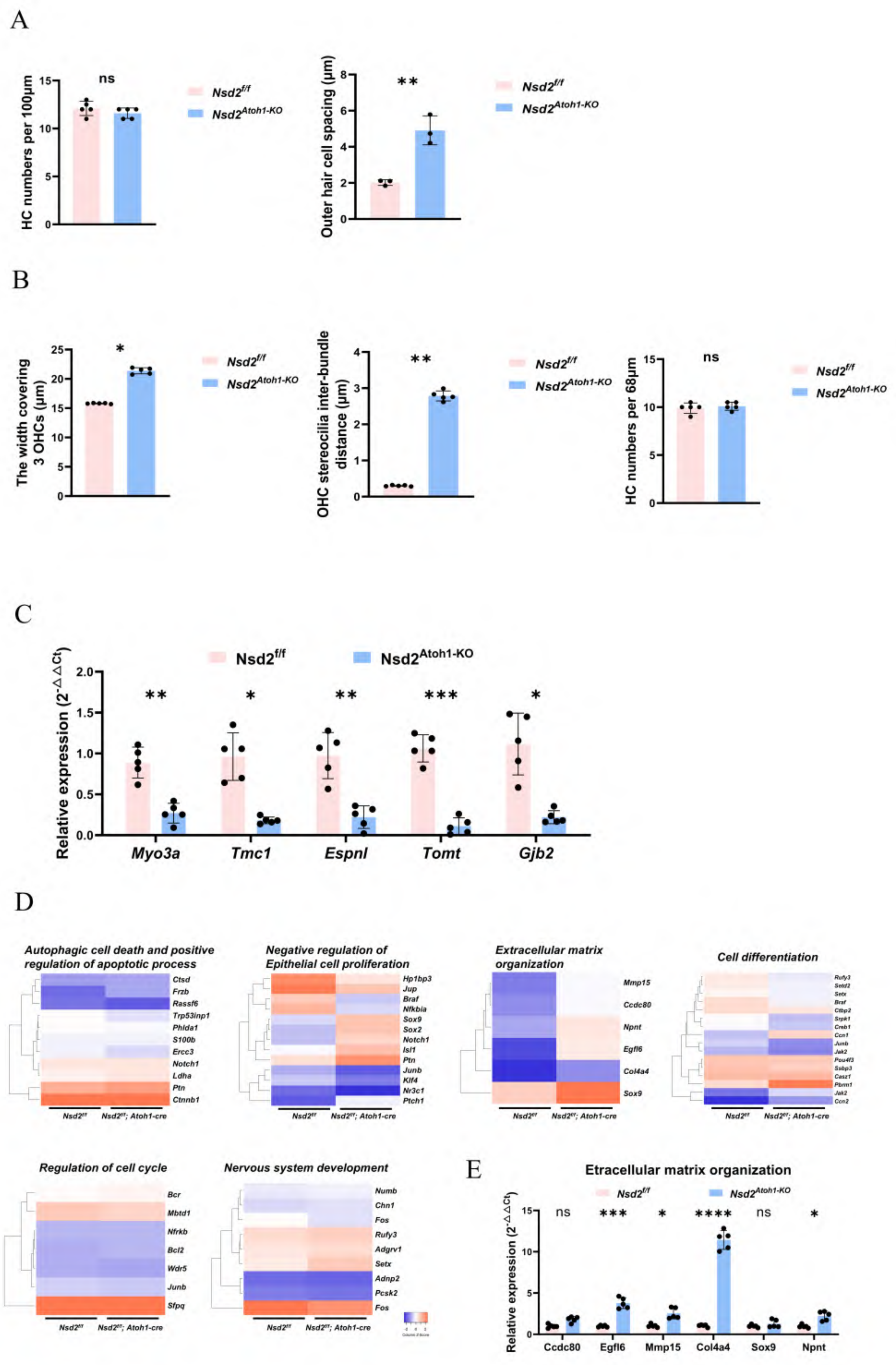
A
HC numbers per 100μm
ns
Nsd2^f/f
Nsd2^Atoh1-KO
Outer hair cell spacing (μm)
**
B
The width covering 3 OHCs (μm)
*
OHC stereocilia inter-bundle distance (μm)
**
HC numbers per 68μm
ns
C
Nsd2^f/f
Nsd2^Atoh1-KO
Relative expression (2^-ΔΔCt)
** * ** *** *
Myo3a
Tmc1
Espnl
Tomt
Gjb2
D
Autophagic cell death and positive regulation of apoptotic process
Ctsd
Frzb
Rassf6
Trp53inp1
Phlda1
S100b
Ercc3
Notch1
Ldha
Ptn
Ctnnb1
Negative regulation of Epithelial cell proliferation
Hp1bp3
Jup
Braf
Nfkbia
Sox9
Sox2
Notch1
Isl1
Ptn
Junb
Klf4
Nr3c1
Ptch1
Extracellular matrix organization
Mmp15
Ccdc80
Npnt
Egfl6
Col4a4
Sox9
Cell differentiation
Regulation of cell cycle
Bcr
Mbtd1
Nfrkb
Bcl2
Wdr5
Junb
Sfpq
Nervous system development
Numb
Chn1
Fos
Rufy3
Adgrv1
Setx
Adnp2
Pcsk2
Fos
Nsd2^f/f
Nsd2^f/f; Atoh1-cre
E
Etracellular matrix organization
Nsd2^f/f
Nsd2^Atoh1-KO
Relative expression (2^-ΔΔCt)
ns *** * **** ns *
Ccdc80
Egfl6
Mmp15
Col4a4
Sox9
Npnt

Fig.S3

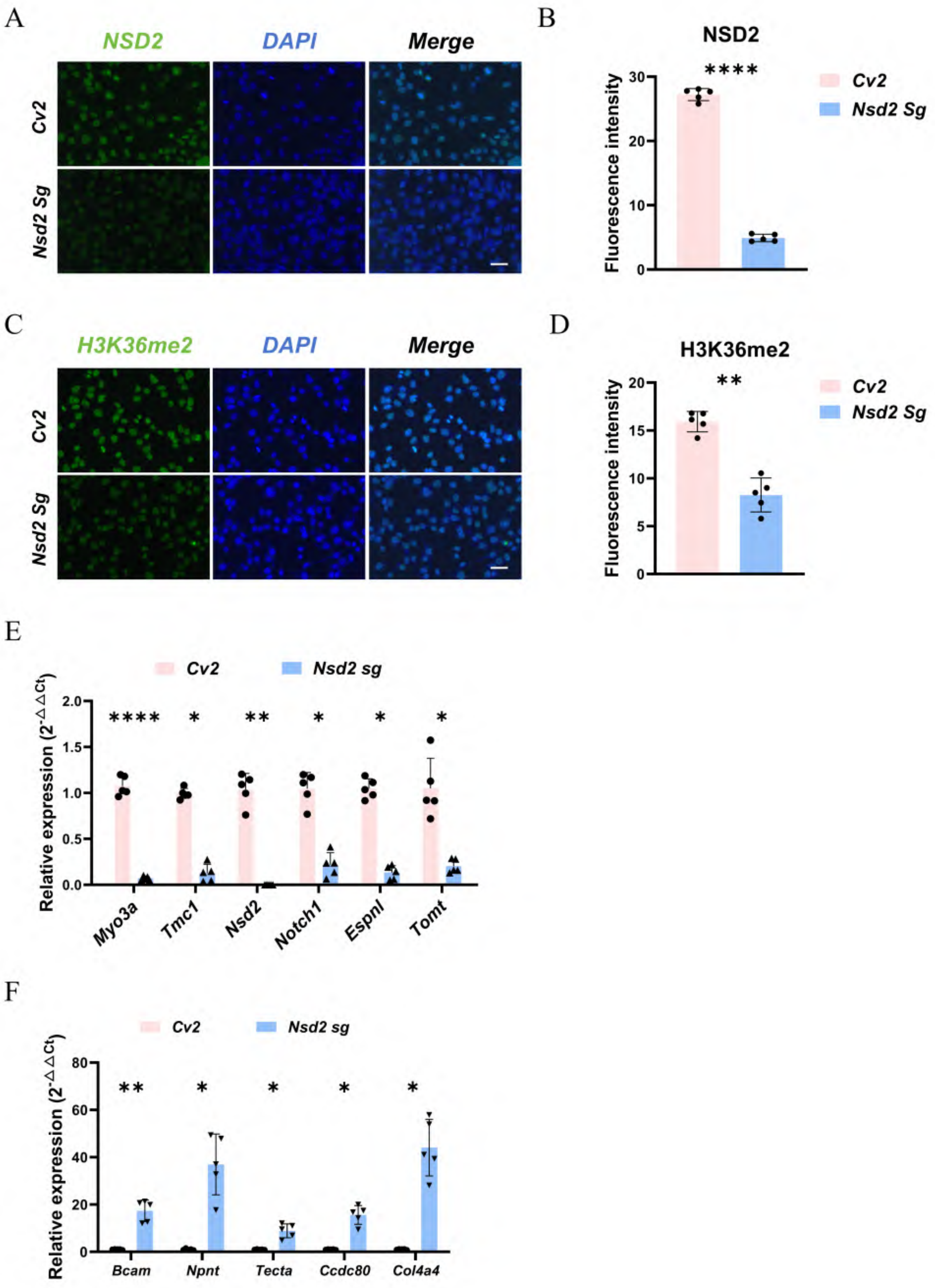
A
NSD2
DAPI
Merge
Cv2
Nsd2 Sg
B
NSD2
****
Cv2
Nsd2 Sg
Fluorescence intensity
0
10
20
30
C
H3K36me2
DAPI
Merge
Cv2
Nsd2 Sg
D
H3K36me2
**
Cv2
Nsd2 Sg
Fluorescence intensity
0
5
10
15
20
E
Cv2
Nsd2 sg
Relative expression ($2^{-\Delta\Delta Ct}$)
0.0
0.5
1.0
1.5
2.0
****
*
**
*
*
*
Myo3a
Tmc1
Nsd2
Notch1
Espnl
Tomt
F
Cv2
Nsd2 sg
Relative expression ($2^{-\Delta\Delta Ct}$)
0
20
40
60
80
**
*
*
*
*
Bcam
Npnt
Tecta
Ccdc80
Col4a4

Fig.S4

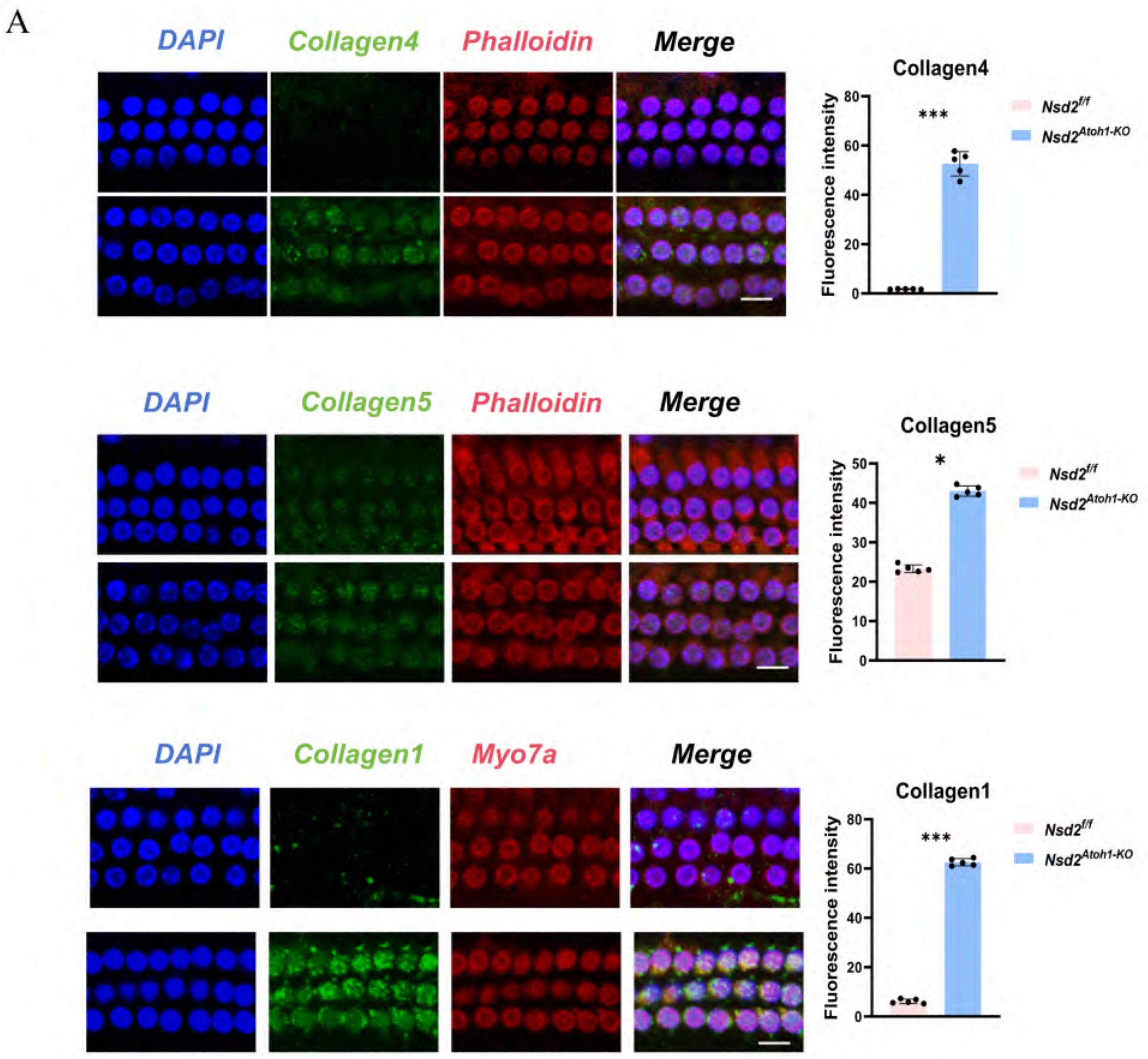
A
DAPI
Collagen4
Phalloidin
Merge
Collagen4
Fluorescence intensity
***
Nsd2f/f
Nsd2Atoh1-KO
DAPI
Collagen5
Phalloidin
Merge
Collagen5
Fluorescence intensity
*
Nsd2f/f
Nsd2Atoh1-KO
DAPI
Collagen1
Myo7a
Merge
Collagen1
Fluorescence intensity
***
Nsd2f/f
Nsd2Atoh1-KO

Fig.S5

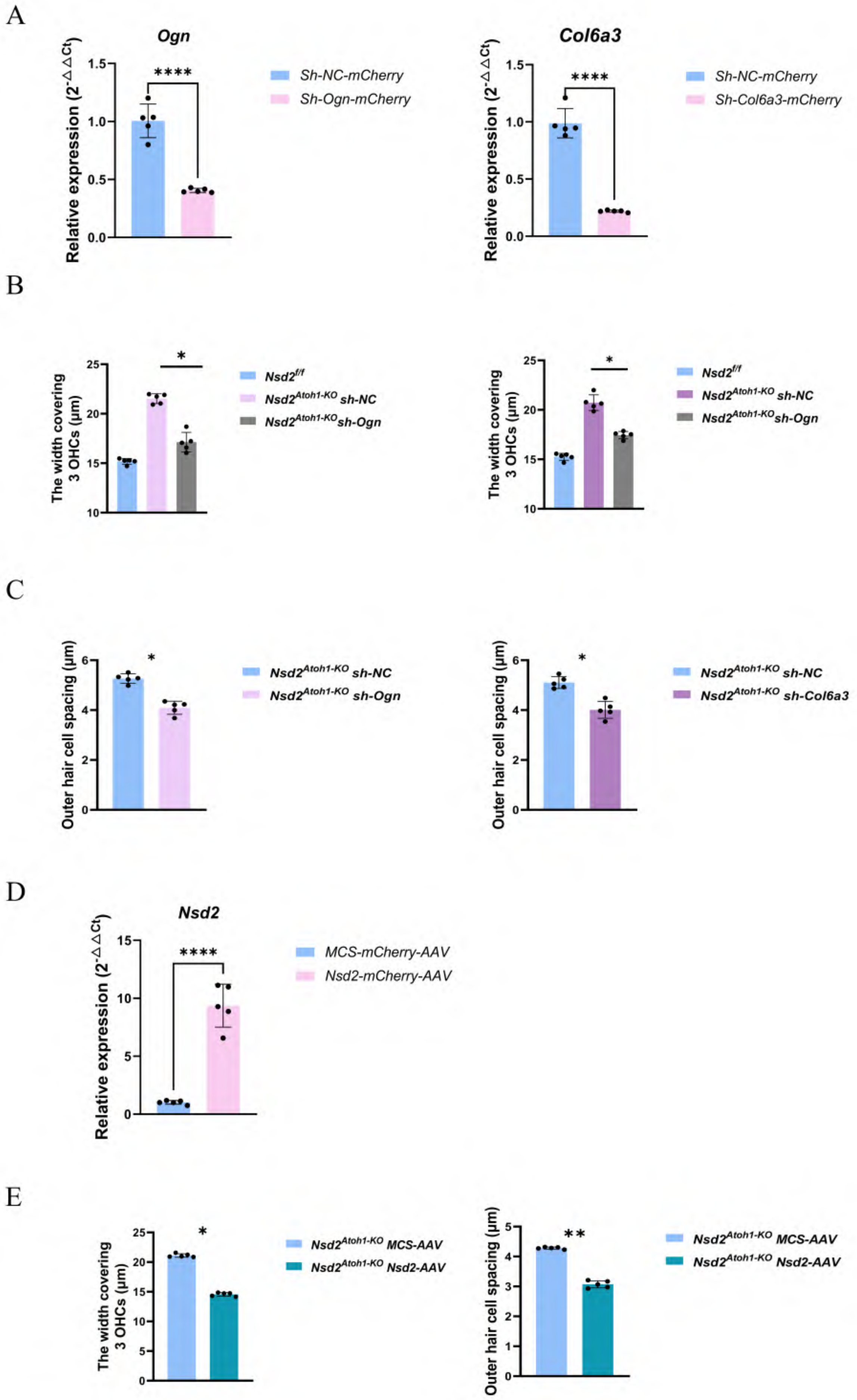
A
Ogn
Relative expression (2^-ΔΔCt)
1.5
1.0
0.5
0.0
****
Sh-NC-mCherry
Sh-Ogn-mCherry
Col6a3
****
Sh-NC-mCherry
Sh-Col6a3-mCherry
B
The width covering 3 OHCs (μm)
25
20
15
10
*
Nsd2f/f
Nsd2Atoh1-KO sh-NC
Nsd2Atoh1-KOsh-Ogn
C
Outer hair cell spacing (μm)
6
4
2
0
*
Nsd2Atoh1-KO sh-NC
Nsd2Atoh1-KO sh-Ogn
Nsd2Atoh1-KO sh-Col6a3
D
Nsd2
Relative expression (2^-ΔΔCt)
15
10
5
0
****
MCS-mCherry-AAV
Nsd2-mCherry-AAV
E
The width covering 3 OHCs (μm)
*
Nsd2Atoh1-KO MCS-AAV
Nsd2Atoh1-KO Nsd2-AAV
Outer hair cell spacing (μm)
**

**Fig.S6** NSD2-deficiency impairs cellular durotaxis on a normal stiffness substrate.

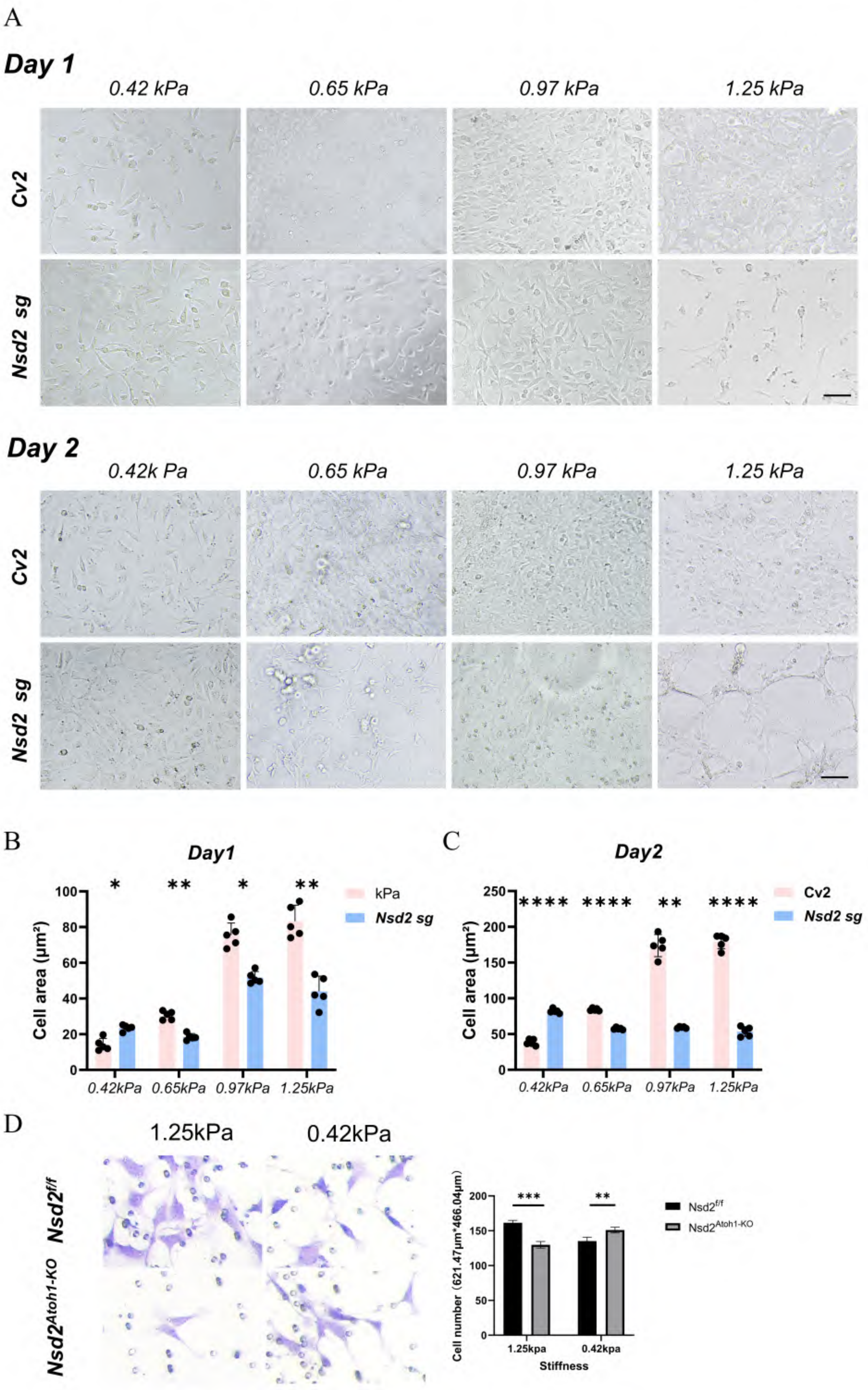

Fig.S7

A

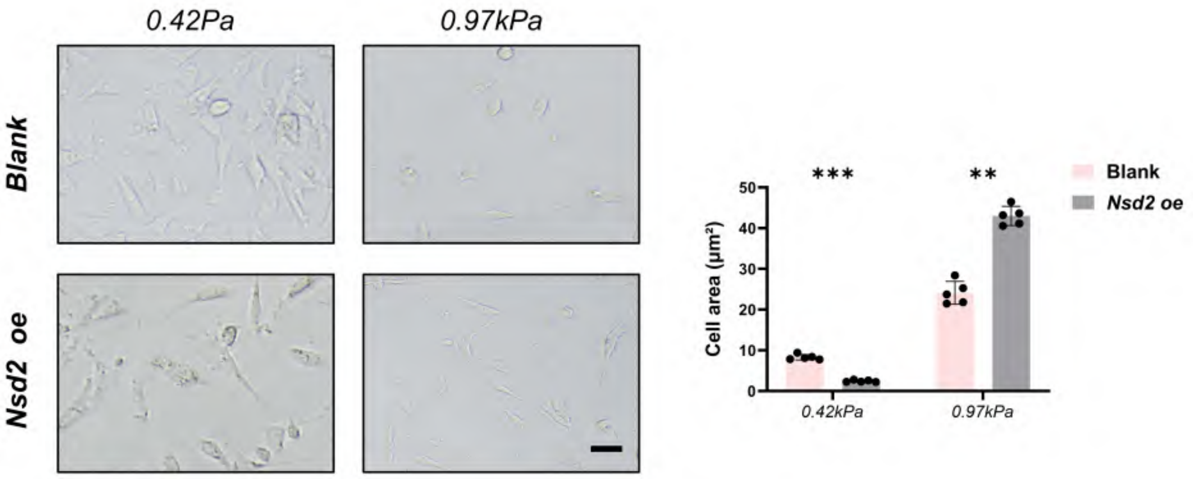

0.42Pa
0.97kPa
Blank
Nsd2 oe
***
**
Blank
Nsd2 oe
Cell area (μm²)
50
40
30
20
10
0
0.42kPa
0.97kPa


B

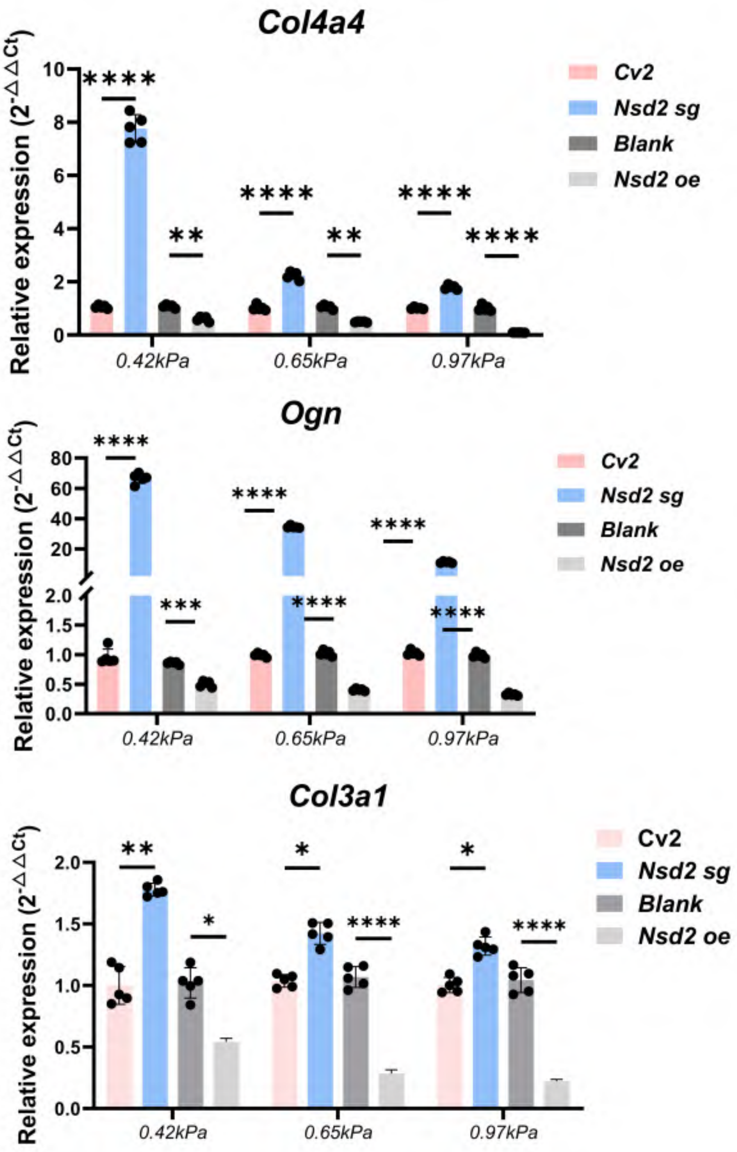

Col4a4
Relative expression (2-ΔΔCt)
10
8
6
4
2
0
****
**
****
**
****
****
0.42kPa
0.65kPa
0.97kPa
Cv2
Nsd2 sg
Blank
Nsd2 oe
Ogn
80
60
40
20
2.0
1.5
1.0
0.5
0.0
****
***
****
****
****
****
Col3a1
2.0
1.5
1.0
0.5
0.0
**
*
*
****
*
****

Fig.S8

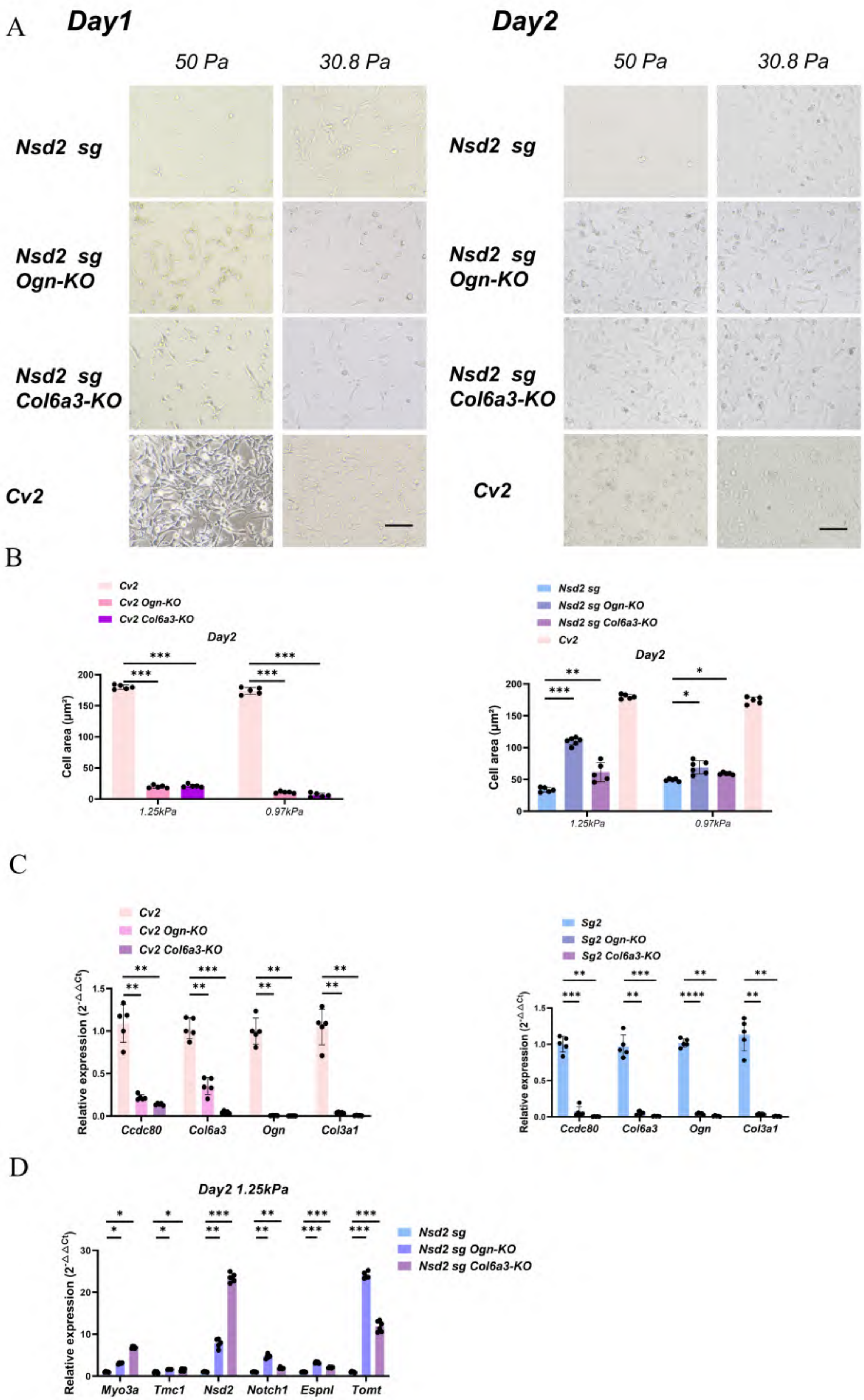

A
Day1
Day2
50 Pa
30.8 Pa
Nsd2 sg
Nsd2 sg Ogn-KO
Nsd2 sg Col6a3-KO
Cv2
B
Cv2
Cv2 Ogn-KO
Cv2 Col6a3-KO
Day2
Cell area (μm²)
1.25kPa
0.97kPa
Nsd2 sg
Nsd2 sg Ogn-KO
Nsd2 sg Col6a3-KO
Cv2
C
Cv2
Cv2 Ogn-KO
Cv2 Col6a3-KO
Relative expression (2^-ΔΔCt)
Ccdc80
Col6a3
Ogn
Col3a1
Sg2
Sg2 Ogn-KO
Sg2 Col6a3-KO
D
Day2 1.25kPa
Nsd2 sg
Nsd2 sg Ogn-KO
Nsd2 sg Col6a3-KO
Myo3a
Tmc1
Nsd2
Notch1
Espnl
Tomt

Fig.S9

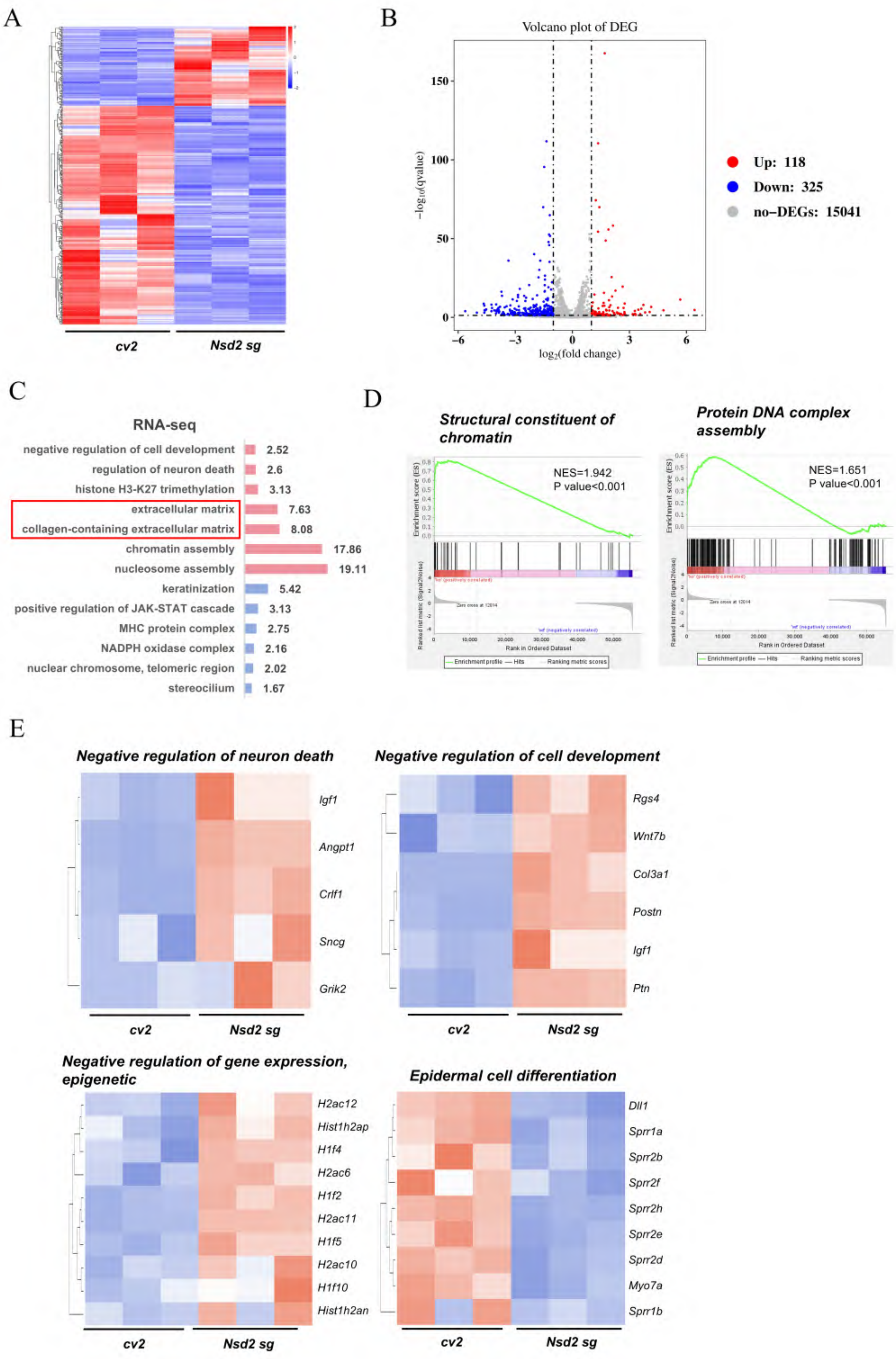
A
cv2
Nsd2 sg
B
Volcano plot of DEG
−log10(qvalue)
log2(fold change)
Up: 118
Down: 325
no−DEGs: 15041
C
RNA-seq
negative regulation of cell development 2.52
regulation of neuron death 2.6
histone H3-K27 trimethylation 3.13
extracellular matrix 7.63
collagen-containing extracellular matrix 8.08
chromatin assembly 17.86
nucleosome assembly 19.11
keratinization 5.42
positive regulation of JAK-STAT cascade 3.13
MHC protein complex 2.75
NADPH oxidase complex 2.16
nuclear chromosome, telomeric region 2.02
stereocilium 1.67
D
Structural constituent of chromatin
NES=1.942
P value<0.001
Protein DNA complex assembly
NES=1.651
P value<0.001
Rank in Ordered Dataset
Enrichment profile — Hits Ranking metric scores
E
Negative regulation of neuron death
Igf1
Angpt1
Crlf1
Sncg
Grik2
cv2
Nsd2 sg
Negative regulation of cell development
Rgs4
Wnt7b
Col3a1
Postn
Igf1
Ptn
cv2
Nsd2 sg
Negative regulation of gene expression, epigenetic
H2ac12
Hist1h2ap
H1f4
H2ac6
H1f2
H2ac11
H1f5
H2ac10
H1f10
Hist1h2an
cv2
Nsd2 sg
Epidermal cell differentiation
Dll1
Sprr1a
Sprr2b
Sprr2f
Sprr2h
Sprr2e
Sprr2d
Myo7a
Sprr1b
cv2
Nsd2 sg

### Fig.S10

A

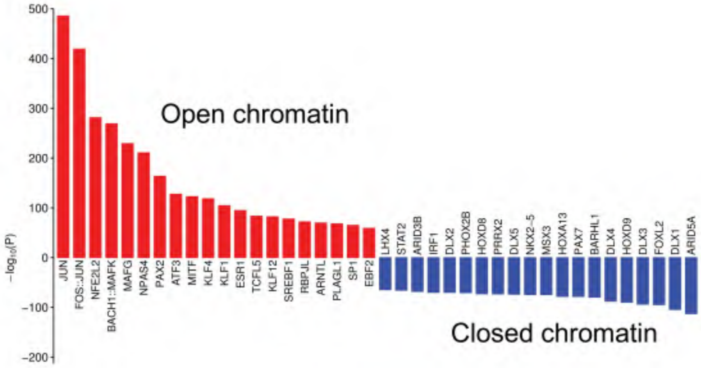
Open chromatin
Closed chromatin
−log10(P)
JUN
FOS::JUN
NFE2L2
BACH1::MAFK
MAFG
NPAS4
PAX2
ATF3
MITF
KLF4
KLF1
ESR1
TCFL5
KLF12
SREBF1
RBPJL
ARNTL
PLAGL1
SP1
EBF2
LHX4
STAT2
ARID3B
IRF1
DLX2
PHOX2B
HOXD8
PRRX2
DLX5
NKX2−5
MSX3
HOXA13
PAX7
BARHL1
DLX4
HOXD9
DLX3
FOXL2
DLX1
ARID5A

B

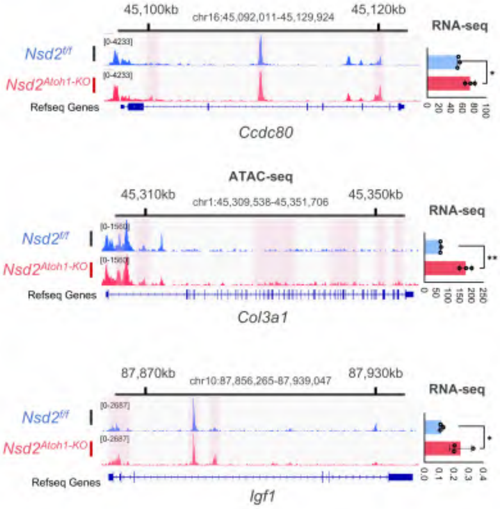
chr16:45,092,011-45,129,924
45,100kb
45,120kb
RNA-seq
Nsd2f/f
Nsd2Atoh1-KO
Refseq Genes
Ccdc80
ATAC-seq
chr1:45,309,538-45,351,706
45,310kb
45,350kb
Col3a1
chr10:87,856,265-87,939,047
87,870kb
87,930kb
Igf1
